# Paternally-acting canonical RNA-directed DNA methylation pathway genes sensitize Arabidopsis endosperm to paternal genome dosage

**DOI:** 10.1101/527853

**Authors:** P. R. V. Satyaki, Mary Gehring

## Abstract

Seed development is sensitive to parental dosage, with excess maternal or paternal genomes creating reciprocal phenotypes. Paternal genomic excess frequently results in extensive endosperm proliferation without cellularization and seed abortion. We previously showed that loss of the RNA Pol IV gene *nrpd1* in tetraploid fathers represses seed abortion in paternal excess crosses. Here we show genetically that RNA-directed DNA methylation (RdDM) pathway activity in the paternal parent is sufficient to determine the viability of paternal excess seeds. We compared transcriptomes, DNA methylation, and small RNAs from endosperm of balanced crosses (diploid × diploid) and lethal (diploid × tetraploid) and viable paternal excess (diploid × tetraploid *nrpd1*). Endosperm from both lethal and viable paternal excess seeds share widespread transcriptional and DNA methylation changes at genes and TEs. Interploidy seed abortion is thus unlikely to be caused by either transposable element or imprinted gene mis-regulation, and its repression by loss of paternal RdDM is associated with only modest gene expression changes. Finally, using allele-specific transcription data, we present evidence for a transcriptional buffering system that increases expression of maternal alleles and represses paternal alleles in response to excess paternal genomic dosage. These findings prompt reconsideration of models for dosage sensitivity in endosperm.

## Introduction

The endosperm of flowering plants is an essential tissue for seed viability. It is most commonly triploid, formed when the diploid central cell is fertilized by a haploid sperm. In many flowering plants, endosperm proceeds through the early phases of proliferative development as a syncytium before differentiating into three sub-types: micropylar, peripheral and chalazal endosperm (Li and Berger, 2012). Cellularization is a critical step in endosperm development, after which cell division slows and eventually ceases (Hehenberger et al., 2012; Li and Berger, 2012). In some species, like Arabidopsis, the endosperm is almost completely degraded at seed maturation as nutrients are assimilated and stored in the embryo (Li and Berger, 2012), whereas in other species, like grasses, it is persistent and mobilized later during seed germination. The endosperm serves several important functions: it mediates resource transfer from the mother to the growing embryo or germinating seedling (Li and Berger, 2012), signals to the maternal integuments to promote their proliferation and allow accommodation of the growing offspring (Figueiredo et al., 2016), is required for embryonic development, and influences seed dormancy and germination (Fiume and Fletcher, 2012; Piskurewicz et al., 2016; Yan et al., 2014). Through these activities, endosperm influences seed size.

Balance between the maternal and paternal genomes in endosperm is important for normal endosperm development and, consequently, seed development and viability. Violations of the 2:1 maternal:paternal (m:p) genome ratio in endosperm lead to developmental defects, although species vary in the extent to which they tolerate violations of this ratio (Cooper, 1951; Esen and Soost, 1973; Håkansson and Ellerström, 1950; Milbocker and Sink, 1969; Muntzing, 1936; Povilus et al., 2018; Scott et al., 1998, 1998; Stoute et al., 2012). In Arabidopsis, increased maternal genome dosage (maternal excess) leads to premature cellularization of the endosperm and the formation of smaller seeds (Scott et al., 1998). By contrast, increased paternal genome dosage (paternal excess) in crosses between diploid mothers and hexaploid fathers leads to a failure of endosperm cellularization, prolonged cell proliferation, and seed abortion (Scott et al., 1998). However, there is intra-specific variation in the levels of seed abortion observed in paternal excess crosses. For example, whereas tetraploid Col-0 induces seed abortion when pollinating diploid mothers, tetraploid C24 and Cvi do not (Dilkes et al., 2008; Lu et al., 2012; Piskurewicz et al., 2016; Scott et al., 1998).

What determines the threshold between seed lethality and viability in conditions of paternal genomic excess? Mutants that repress seed abortion in interploidy crosses, as well data from incompatible interspecific crosses, which share features with interploidy crosses, have provided some answers to this question. Components in the endosperm, in the gametophytes, or in the parental sporophytes, have been proposed to be responsible for endosperm dosage sensitivity in interploid crosses. Some of the elements that determine the critical threshold might be linked to interactions between the endosperm and maternal genotype. In one model, the imbalance is between the paternal dose and the female gametophyte (Birchler, 2014; von Wangenheim and Peterson, 2004). A second model involves interactions between the paternal excess endosperm and the diploid maternal integuments, which develop into the seed coat (Muntzing, 1936). The ability of maternal sporophytic mutations in the flavonoid pathway to repress paternal excess seed abortion is consistent with this model (Dilkes et al., 2008; Doughty et al., 2014). The levels of *AGLs*, the small RNA *siRNA854*, chromatin regulators, cell wall genes, and defense response genes, all presumed to be acting in the endosperm, have been linked to paternal excess seed abortion (Burkart-Waco et al., 2013; Jiang et al., 2017; Kradolfer et al., 2013; Walia et al., 2009; Wang et al., 2018; Wolff et al., 2015).

Multiple models suggest that endosperm ploidy incompatibilities are caused by epigenetic abnormalities. The epigenetic mark DNA methylation is established and partially maintained through the RNA-directed DNA methylation (RdDM) pathway. During this process, short non-coding RNAs generated by RNA Pol IV are converted into double-stranded RNAs by RDR2. These are then subsequently diced by DCL3 into 24 nt small RNAs (sRNAs), loaded into an ARGONAUTE complex (usually AGO4 or AGO6), and then interact with a non-coding RNA transcribed by another polymerase, RNA Pol V, to direct DNA methylation by the *de novo* methyltransferase DRM2. In normal triploid endosperm, maternal chromosomes are DNA hypomethylated relative to paternal chromosomes due to the activity of the DNA demethylase *DME* (Gehring et al., 2009; Hsieh et al., 2009; Ibarra et al., 2012). DNA methylation represses expression of some endosperm genes and promotes the expression of others (Satyaki and Gehring, 2017). It has been proposed that a cause of interploid seed abortion is increased expression of transposable elements (TEs) due to epigenetic alterations (Martienssen, 2010). Under this model, a maternally deposited sRNA dose is insufficient to silence the doubled number of paternally inherited TEs. Another model is influenced by the parental conflict theory (Haig and Westboy, 1991). The theory argues that in a polyandrous system in which the mother provisions resources for her offspring from various fathers, genes restricting resource allocation to any one seed and favoring equitable distribution of resources across all the progeny become predominantly maternal in expression (maternally expressed genes or MEGs). On the other hand, genes promoting resource allocation and larger seed production are predominantly paternally expressed (paternally expressed genes or PEGs). This type of allele-specific expression is an epigenetic phenomenon referred to as gene imprinting. According to these ideas, an excess dose of paternal chromosomes leads to over-expression of PEGs (genes that promote endosperm proliferation) eventually leading to seed abortion. Consistent with this model, PEGs were reported to be over-expressed in paternal excess seeds (Kradolfer et al., 2013; Schatlowski et al., 2014) and loss-of-function mutations in multiple PEGs repress seed abortion in conditions of paternal genomic excess (Huang et al., 2017; Jiang et al., 2017; Kradolfer et al., 2013; Wolff et al., 2015).

We recently proposed another model based on discoveries about the function of *NRPD1*, which codes for the largest subunit of RNA Pol IV, in normal triploid endosperm (Erdmann et al., 2017). Expression of 24 nt sRNAs in *Arabidopsis thaliana* endosperm is paternally biased and concentrated in pericentromeric heterochromatin, but a subset of 24 nt sRNAs are expressed predominantly from the maternal alleles of genes (Erdmann et al., 2017). We showed that paternal loss of *NRPD1* in 2m:1p triploid endosperm increased the maternal fraction of the transcriptome, suggesting that wild-type *NRPD1* represses maternal genome dosage, probably via the production of 24 nt small RNAs. We also discovered that paternally inherited mutations in *NRPD1* repress seed abortion caused by excess paternal genomes (i.e. 2N Col × 4N Col), leading to the hypothesis that the increased maternal fraction of the transcriptome in *nrpd1* mutant endosperm compensates for increased paternal genomic dosage. In this model, the loss of RNA Pol IV-dependent sRNA production in the endosperm is essential for viability (Erdmann et al., 2017).

Finally, the “easiRNA” model argues that *NRPD1* and *RDR6* act together in a non-canonical pathway in the male gametophyte to produce easiRNAs whose concentration scales with ploidy (Borges et al., 2018; Martinez et al., 2018). easiRNAs are defined as 21-22 nt sRNAs produced from the processing of epigenetically activated TE mRNAs (Creasey et al., 2014). It is proposed that easiRNAs are transmitted during fertilization from sperm to the central cell, where they inhibit RdDM in the developing endosperm (Borges et al., 2018; Martinez et al., 2018). In this model, easiRNAs are lost from sperm when the paternal copy of *NRPD1* is mutated. The loss of paternal easiRNAs allows the restoration of a functional RdDM pathway in endosperm using maternal copies of *NRPD1*. The restored RdDM pathway was proposed to repress excess paternal dosage and restore seed viability.

To test among these models and further understand the nature of intolerance or tolerance to paternal genomic excess, we tested the genetic and molecular contributions to interploidy seed abortion and repression. We created tetraploid mutants for members of the canonical and non-canonical small RNA pathways to test the genetic requirements for seed abortion in paternal excess crosses. We also profiled sRNAs, DNA methylation, and gene expression in balanced (2N × 2N), lethal paternal excess (2N × 4N), and viable paternal excess (2N × 4N *nrpd1*) endosperm. We found that genes of the canonical RdDM pathway were necessary in the male parent only for seed abortion induced by paternal genomic excess. Through transcriptomic profiling we found that there were extensive gene and transposable element expression changes associated with lethal paternal genomic excess but that only a small fraction were ameliorated in viable paternal excess endosperm. Our analysis also revealed the signatures of a potential buffering system that attempts to rebalance transcription in conditions of paternal excess by repressing paternal alleles and activating maternal alleles. These observations indicate that endosperm development can tolerate surprisingly large variations in gene expression at most genes and suggests that relatively small gene expression changes contribute to the critical dosage that determines seed viability under conditions of paternal genomic excess.

## Results

### Paternal loss of canonical RdDM pathway genes is sufficient for suppression of interploid seed abortion

In crosses between diploid females and tetraploid males, tetraploid *nrpd1* mutant fathers repress seed abortion, whereas diploid *nrpd1* mothers have no effect (Erdmann et al., 2017; Martinez et al., 2018). We tested if other members of the canonical RdDM pathway behaved genetically in the same manner. Using colchicine-induced tetraploid mutants of *shh1, rdr2, dcl3, nrpe1*, and *drm2*, we examined seeds obtained from crosses between 2N wild-type (Col) mothers and 4N (Col) fathers that were either wild-type or mutant for one of the RdDM pathway components (Figure 1). Mature seeds were scored as normal, aborted, or abnormal and multiple independent crosses were analyzed. Typically, seed abortion in each set of crosses varied by between 20 and 30%, regardless of whether parents were wild-type or mutant (Figure 1A). For example, in wild-type paternal excess crosses (2N Col × 4N Col) seed abortion ranged from 57.8%-100%, with mean seed abortion at 81.1%. Therefore, we considered only those mutants capable of substantially enhancing seed viability across multiple crosses as being true repressors of seed abortion.

**Figure 1:**
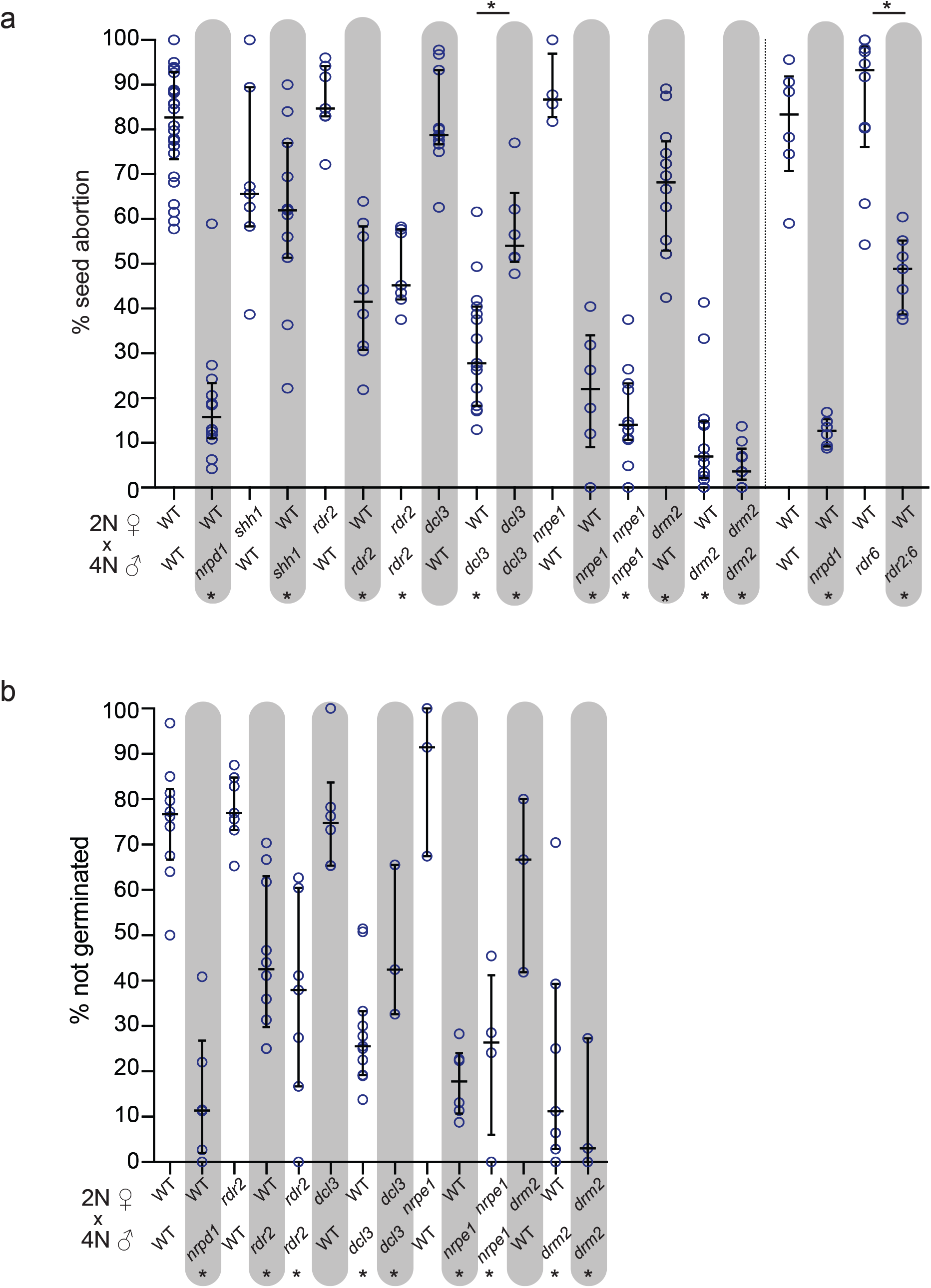
Loss of paternal RdDM genes but not *RDR6* represses seed abortion in paternal excess crosses. **a)** Each circle represents seed abortion rate in one cross and represents multiple siliques from a single inflorescence. **b)** Each circle represents the percent of seeds that failed to germinate from each scored collection of seed. Failure to germinate was defined as the inability to produce either a radicle or a hypocotyl. Bars show median and interquartile range. * at bottom represents statistically significant difference (p<0.05) in comparisons between indicated cross and cross between wild-type (WT) diploid (2N) Col-0 mothers and wild-type tetraploid (4N) Col-0 fathers. * at top represents statistically significant differences (p<0.05) between crosses indicated. Statistical significance calculated by Wilcoxon test.

All paternally inherited RdDM pathway mutations, except *shh1*, substantially repressed seed abortion and promoted the production of fully developed seeds capable of germination (Figure 1, Supplemental Figure 1). SHH1 is required for RNA Pol IV recruitment at a subset of its target sites (Law et al., 2013), suggesting that SHH1-independent activity of RNA Pol IV is important for seed abortion. Mutations in *nrpe1* (encoding the largest subunit of RNA Pol V) and *drm2* mirrored *nrpd1* in their substantial repression of seed abortion; mean seed abortion among examined seeds was 18.9% for *nrpd1*, 21.4% for *nrpe1* and 11.3% for *drm2* (Figure 1A, Supplemental Figure 1). *rdr2* and *dcl3* mutations repressed seed abortion to a lesser extent (Figure 1, Supplemental Figure 1). The previously described redundancy between *DCL3* and other *DCL* paralogs (Gasciolli et al., 2005; Stroud et al., 2013) likely explains the lower repression of seed abortion by loss of *DCL3*, but it remains unclear as to why loss of *RDR2* does not repress seed abortion to the same extent as that observed upon loss of *NRPD1*.

For all mutants the seed abortion ratios were reflected in the percentage of seeds that germinated on plates (Figure 1B, Supplemental Figure 1). Tetraploid *drm2* and *nrpd1* mutants also suppressed interploidy seed abortion when crossed with wild-type diploid L*er* mothers (Supplemental Figure 1E,G) (Erdmann et al., 2017), indicating the effect was not specific to Col mothers. RNA Pol IV has also been proposed to function in partnership with RDR6 in a non-canonical RdDM pathway (Martinez et al., 2018). We therefore tested if tetraploid fathers mutant for *rdr6* could suppress paternal excess seed abortion. A paternally inherited *rdr6* mutation did not repress seed abortion and did not act additively with *rdr2* (Figure 1A, Supplemental Figure 1B). We also tested whether mutations in the CHG methyltransferase *CMT3* could suppress interploidy seed abortion when inherited through the male – no effect was observed (Supplemental Figure 1D,F).

In contrast to repression of seed abortion by paternally inherited loss-of-function mutations in RdDM pathway genes, most loss-of-function mutations inherited through the diploid mother did not repress seed abortion in paternal excess crosses with wild-type tetraploid fathers (Figure 1, Supplemental Figure 1A,C). An exception was maternal loss of *DRM2*, which resulted in a statistically significant repression in seed abortion, although the magnitude of the repression was small (mean percentage of aborted seed was 66.9% for *drm2* compared to 81.1% for WT) and did not phenocopy the extensive reduction in seed abortion observed upon the loss of paternal RdDM pathway components (Figure 1, Supplemental Figure 1A,C). It was previously reported that suppression of paternal excess seed abortion also occurs when both parents are mutant for *NRPD1* (Erdmann et al., 2017; Martinez et al., 2018). We observed the same effect for all other genes tested: *rdr2*, *dcl3*, *nrpe1*, and *drm2* (Figure 1, Supplemental Figure 1A,C). In sum, these genetic results indicate 1) that paternal loss of the canonical RdDM pathway is sufficient to suppress seed abortion caused by paternal genomic excess and 2) that a genetically complete canonical RdDM pathway is not required in endosperm itself for suppression of seed abortion.

### Massive, shared gene mis-regulation in lethal and viable paternal excess endosperm

To determine what genes or processes were associated with interploidy seed lethality and its genetic suppression, we examined the transcriptional effects of doubling paternal dosage in the endosperm. We performed mRNA-seq to high depth on developing endosperm from three biological replicates of balanced crosses (L*er* × Col), lethal paternal excess (L*er* × 4N Col), and viable paternal excess (L*er* × 4N Col *nrpd1*) (Figure 2, Supplemental Table 1). Surprisingly, comparisons of the three transcriptomes by principal component analysis indicated that lethal and viable paternal excess endosperm were more similar to each other than to balanced endosperm (Figure 2A). About a third of the transcriptome was significantly mis-expressed (2-fold or greater difference at q < 0.05) in lethal paternal-excess endosperm compared to balanced endosperm: 4,054 genes were more highly expressed and 3,855 genes decreased in expression (Figure 2B, Supplemental Dataset 1). GO analyses (Supplemental Dataset showed that genes with lower transcript abundance in lethal paternal excess relative to balanced endosperm were enriched for those encoding light harvesting proteins, proteins in glucose and starch metabolism, hormone responses, response to abiotic stimuli, and cell wall organization. Whereas the predicted consequences of many of these changes remain unclear, the decreased expression of cell wall genes is consistent with the failure in endosperm cellularization that has been previously described in lethal paternal excess crosses (Wolff et al., 2015). GO analyses of genes with increased expression identified enrichment for those encoding proteins involved in protein deneddylation, ribosome biogenesis, DNA replication, chromosome segregation, the cell-cycle. The increased expression of these genes is consistent with increased cell proliferation observed in paternal excess endosperm (Scott et al., 1998; Tiwari et al., 2010). Compared to balanced endosperm, viable paternal excess endosperm also showed a similar quantitative change in gene expression: 3150 genes increased in expression and 2845 decreased (Figure 2B, Supplemental Dataset 1). Many of same genes were mis-regulated in viable and lethal paternal excess endosperm compared to balanced endosperm. Compared to lethal paternal excess, only 188 genes were reduced in viable paternal excess endosperm and 426 genes were more highly expressed (Figure 2B, Supplemental Dataset 1). To further test if the viability of paternal excess seeds was predicated on the transcriptome coming nearer to the dosage of balanced endosperm, we examined the extent to which gene expression was “corrected” in viable paternal excess endosperm (Figure 2C). Most genes with increased expression in lethal paternal excess endosperm were not corrected, but a small subset of genes with decreased expression in lethal paternal excess were moderately corrected in viable paternal excess (Figure 2C).

**Figure 2:**
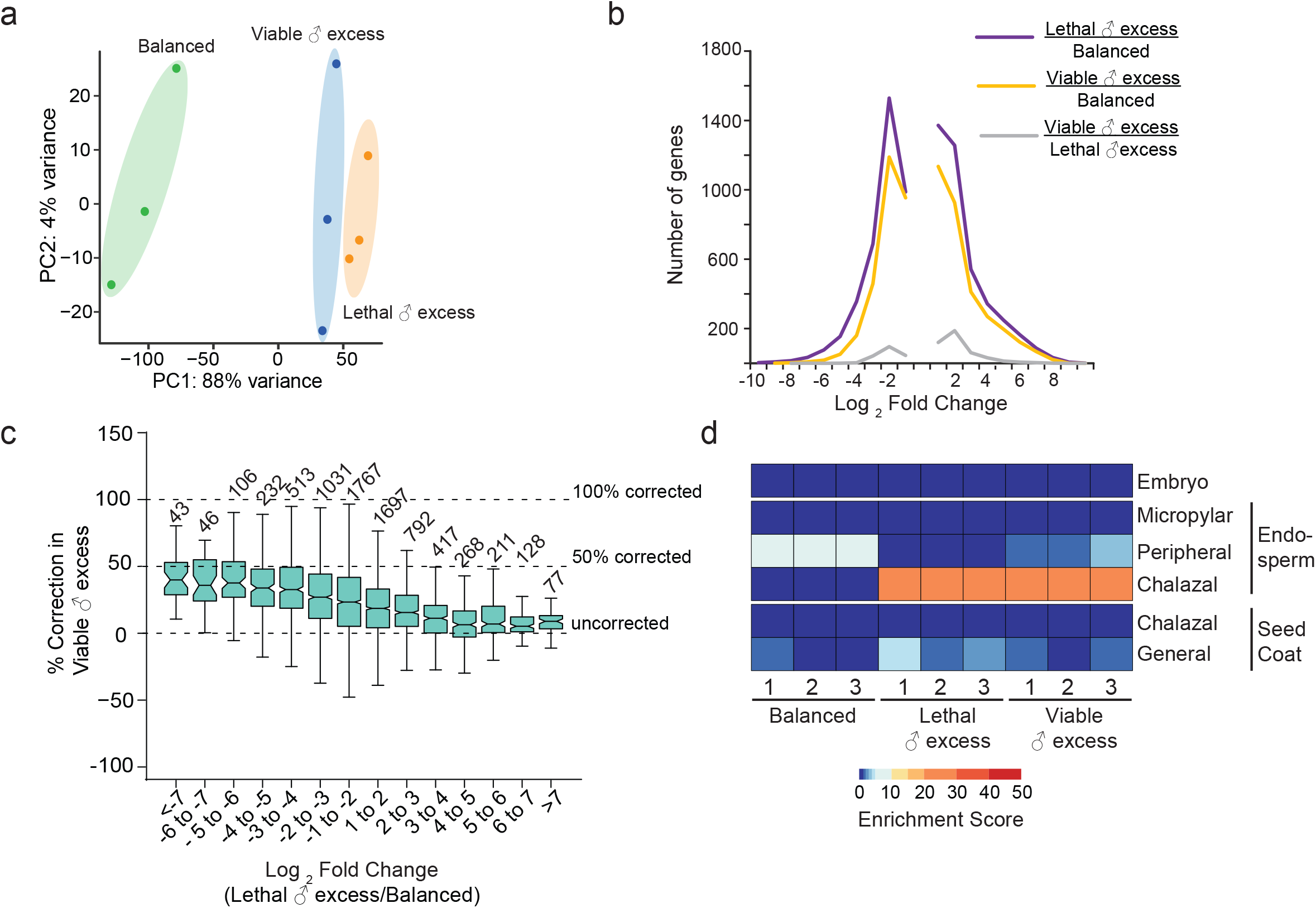
Lethal and viable paternal excess endosperm are transcriptionally more similar to each other than to balanced endosperm. **a)** PCA plot of read counts for genes from biological replicate mRNA-Seq samples. **b)** Plot of number of genes differentially expressed in comparisons of balanced endosperm with both lethal (purple) and viable (yellow) ♂ excess endosperm. Only 614 genes were differentially expressed between viable and lethal ♂ excess endosperm (gray). **c)** Correction value in viable paternal excess endosperm for each gene that was called as being significantly differentially expressed in comparisons of balanced and lethal ♂ excess endosperm. The value was calculated as % Correction= 100-(((log_2_ (viable/balanced)/log_2_ (lethal/balanced))*100). A value of 100% indicates that the gene, which was mis-regulated in lethal ♂ excess, was not differentially expressed in viable ♂ excess relative to balanced endosperm. A value of 0% represents similar mis-regulation in both lethal and viable ♂ excess relative to balanced endosperm. Fold change values and significance for fold-change for b and c were calculated using Cuffdiff. Boxplot is a Tukey plot. **d)** Lethal paternal excess endosperm was enriched for chalazal endosperm gene expression; viable paternal excess endosperm showed both chalazal and peripheral markers. Tissue enrichment for each biological replicate is shown.

We analyzed the transcriptomic data using a tissue enrichment tool to assess if the abundances of markers enriched in chalazal, peripheral, and micropylar endosperm were altered in paternal excess endosperm (Schon and Nodine, 2017). Paternal excess endosperm adopted a gene expression program characteristic of chalazal endosperm (Figure 2D), consistent with phenotypic analysis of lethal paternal excess seeds (Martinez et al., 2018; Scott et al., 1998; Wolff et al., 2015). Viable paternal excess endosperm transcriptomes also showed increased chalazal endosperm marker gene expression, but there was also slightly elevated peripheral endosperm marker gene expression relative to lethal paternal excess endosperm. These results suggest that lethal and viable paternal excess endosperm differ similarly from balanced endosperm in gene expression and subsequent developmental programs.

We also reanalyzed published endosperm transcriptome data from lethal and viable paternal excess endosperm generated using *osd1* and *osd1 nrpd1* mutations (Martinez et al., 2018) (Supplemental Figure 2). Consistent with our data, we found that both L*er* × Col *osd1* (lethal paternal excess) and L*er* × Col *osd1 nrpd1* (viable paternal excess) showed elevated expression of chalazal markers (Supplemental Figure 2D). Endosperm from L*er* × Col *osd1 nrpd1* also showed extensive genic mis-regulation, but the percentage of the transcriptome that was mis-regulated was lower than in our viable paternal excess endosperm datasets (Figure 2B, Supplemental Figure 2B). Additionally, the extent of gene expression correction in viable paternal excess endosperm was also lower in our datasets compared to theirs (Figure 2C, Supplemental Figure 2C). These discrepancies could stem from multiple differences between our experiments. Our endosperm data also had more replicates, higher mappable read depth, and lower levels of seed coat contamination (Supplemental Figure 2, Supplemental Table 1, (Martinez et al., 2018)). We created paternal excess endosperm using tetraploid fathers while Martinez et al. created paternal excess via the use of the *osd1* mutation, which causes omission of the second meiotic division and thus generates diploid pollen. There might be biological differences, as yet unclear, between diploid sperm produced from a diploid parent compared to diploid sperm produced from a tetraploid parent. Also, the *osd1* mutation was backcrossed into Col from another accession (d’Erfurth et al., 2009; Martinez et al., 2018) and interploidy seed abortion is sensitive to the genetic background of both parents (Lu et al., 2012; Piskurewicz et al., 2016; Scott et al., 1998).

### Mis-regulation of imprinted genes characterizes both lethal and viable paternal excess endosperm

Imprinted gene mis-regulation in the endosperm has been suggested as a culprit for the developmental catastrophe of paternal excess crosses (Gutierrez-Marcos et al., 2003; Haig and Westboy, 1991). We found that imprinted genes are disproportionately more likely to show increased expression than all genes in the genome in conditions of lethal paternal excess (N-1 chi-square test, *p= 3×10^−4^* for MEGs and *p<10^−4^* for PEGs) (Figure 3). Of 43 previously identified Col-L*er* PEGs and 130 Col-L*er* MEGs (Pignatta et al., 2014), 25 PEGs (58%) and 27 MEGs (19%) displayed at least a two-fold increase in transcript abundance in lethal paternal excess endosperm (Figure 3A). The abundance of two PEGs and 18 MEGs decreased. The vast majority of these imprinted genes remained mis-regulated in viable paternal excess endosperm (Figure 3B,C). Only two MEGs were differentially expressed in comparisons of viable and lethal paternal excess endosperm – *JLO* transcript abundance increased 2.6-fold and *GSR1* transcript abundance decreased 2.4-fold (Figure 3C, Supplemental Dataset 1). Similarly, only one PEG, AT4G20800, had lower transcript abundance in viable paternal excess endosperm compared to lethal paternal excess endosperm (Figure 3C, Supplemental Dataset 1). These results suggest that mis-regulation of multiple imprinted genes in endosperm is unlikely to be the cause of endosperm dysfunction and seed lethality in interploidy crosses.

**Figure 3:**
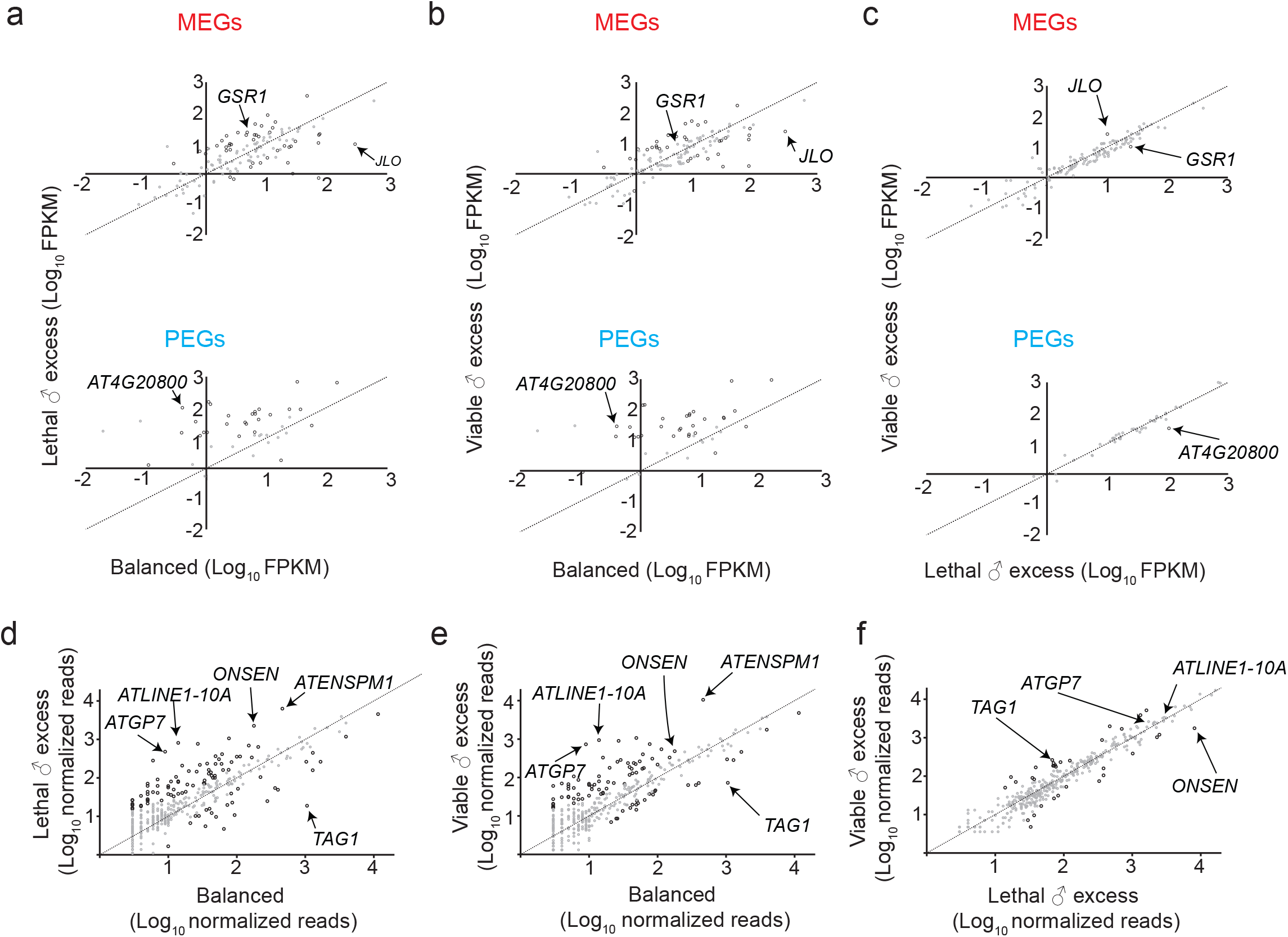
Imprinted genes and transposons are mis-regulated in both lethal and viable ♂ excess endosperm. **a-c)** Expression of Col-L*er* imprinted genes in endosperm. FPKM is normalized expression; statistical significance of difference in abundance calculated by Cuffdiff. q<0.05 represented by black circles. q>0.05 represented by gray circles. **d-f)** Expression from transposable elements is elevated in lethal and viable ♂ excess endosperm. RNA-Seq reads were mapped to consensus sequences from Repbase. Black circles represent TEs with significant differences in transcript abundances according to DEGseq. Gray circles represent TEs without significant differences in transcript abundances.

It was previously demonstrated that mutations in a subset of PEGs or their interactors can suppress paternal excess seed abortion when inherited paternally (Wolff et al., 2015; Huang et al., 2017; Kradolfer et al., 2013; Jiang et al., 2017). We found that whereas several of these genes indeed have increased transcript abundance in lethal paternal excess relative to balanced endosperm, their expression remains altered in viable paternal excess (Supplemental Figure 3). This observation suggests that seed viability brought about by loss of paternal *nrpd1* is independent of gene expression normalization in endosperm of genes whose paternal loss also represses seed abortion.

In summary, these gene expression data indicate that the expression of only a small number of genes distinguishes lethal paternal excess endosperm from viable paternal excess endosperm at this stage of development and suggest that correction of gene expression to levels observed in balanced endosperm, even for imprinted genes, is not necessary for paternal excess seed viability.

### A subset of TEs are mis-regulated in lethal and viable paternal excess seeds

Increased expression of transposable elements (TEs) in paternal excess endosperm has been speculated as a cause of seed abortion (Castillo and Moyle, 2012; Martienssen, 2010). We compared TE expression levels in endosperm by mapping mRNA-seq reads to 375 consensus TE sequences from REPBASE (Figure 3D-F) (Bao et al., 2015). In lethal paternal excess endosperm, 73 families showed a statistically significant change in transcript abundance and 40 increased by 5-fold or more, including the transpositionally active family *ONSEN/ATCOPIA78* (Figure 3D, Supplemental Dataset 3). The TE families *ATLINE1-10A* and *ATGP7* exhibited the largest increases, at least 52-fold (Figure 3D). Additionally, the transcript abundance of 26 TE families decreased in paternal excess crosses. *TAG1*, a transpositionally active TE that is present only in the maternally inherited L*er* genome (Tsay et al., 1993), was the most repressed, nearly 56-fold (Figure 3D). Like for genes, viable paternal excess endosperm displayed a TE expression profile similar to lethal paternal excess endosperm (Figure 3E,F, Supplemental Dataset 3). Only 14 TE families exhibited decreased transcript abundance in viable paternal excess endosperm relative to lethal endosperm and 19 TE families exhibited increased transcript abundance (Figure 3F, Supplemental Dataset 3). The observation that lethal and viable paternal excess endosperm exhibit TE mis-regulation to a similar extent argues that TE mis-regulation by itself is unlikely to be the cause of seed abortion induced by paternal genomic excess.

### The RdDM pathway is attenuated in paternal excess endosperm

A number of proteins that establish or maintain CG, CHG and CHH methylation were differentially expressed among balanced, lethal, and viable paternal excess endosperm (Figure 4, Supplemental Dataset 1). The differential expression of genes encoding members of the RdDM pathway is particularly noteworthy. With the exception of *NRPD1*, the expression of genes encoding subunits of RNA Pol IV and V, along with *RDR2*, *AGO4*, and *DRM2*, was significantly decreased in lethal paternal excess endosperm (Figure 4A, Supplemental Dataset 1). Consistent with these findings, the expression of the 5-methylcytosine DNA glycosylase *ROS1*, whose expression is directly promoted by RdDM (Williams et al., 2015), was also reduced (Figure 4A). The expression of most of these genes increased slightly in viable paternal excess endosperm compared to lethal endosperm, but only increases in *RDR2* and *ROS1* were significant (Figure 4A).

**Figure 4:**
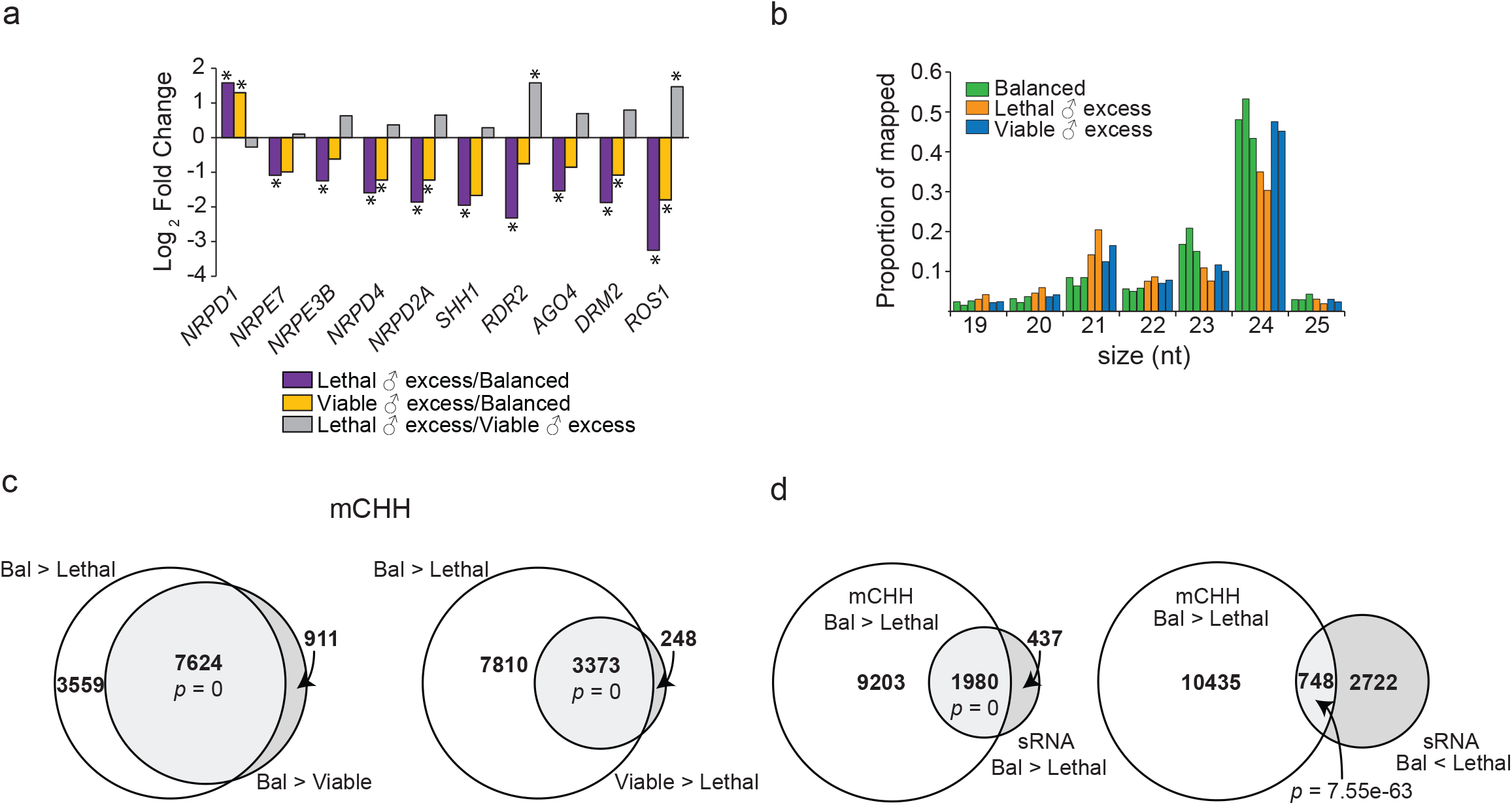
Canonical RdDM pathway function in endosperm is affected by paternal excess. **a)** Genes encoding RdDM components are down-regulated in lethal and viable ♂ excess endosperm. ROS1 expression is a read-out of RdDM activity and reflects differential activity of RdDM. * represents statistically significantly different gene expression. **b)** Small RNA production is impacted in paternal excess endosperm. Size profiles of sRNA reads mapped to TAIR10 genome for three replicates of balanced endosperm and two replicates each of lethal and viable paternal excess endosperm. **c)** CHH methylation losses in paternal excess endosperm. Venn diagram shows intersections of CHH DMRs obtained from comparisons of balanced, lethal, and viable paternal excess endosperm. Lethal and viable paternal excess endosperm share a significant proportion of regions that are hypomethylated relative to balanced endosperm. A smaller subset of DMRs lose more methylation in lethal relative to viable. **d)** Loss of CHH methylation is associated with loss of 24 nt sRNAs. Venn diagrams show the relationship between changes in sRNA abundance and CHH methylation levels. A subset of CHH DMRs are associated with loss of sRNAs. A smaller subset is associated with gains in 24 nt sRNAs. Significance of overlaps was calculated using the Fisher test option from Bedtools.

We tested if sRNA production was impacted in paternal excess endosperm. We sequenced endosperm small RNAs from two replicates of lethal and viable paternal excess endosperm (Supplemental Table 1) and compared them to previously published sRNA data from balanced endosperm (Erdmann et al., 2017). Previous evaluations of small RNAs in paternal excess crosses have been performed on whole seeds, not endosperm, but noted reduced 24 nt sRNAs in lethal paternal excess seeds (Lu et al., 2012; Martinez et al., 2018). We assessed the overall functionality of small RNA production in the endosperm by examining small RNA size profiles. In lethal paternal excess endosperm, 24 nt sRNAs represented the most abundant size class, but their proportion was reduced compared to balanced endosperm (Figure 4B). In viable paternal excess endosperm, the proportion of 24 nt sRNAs was comparable to that of balanced endosperm (Figure 4B). Genic and TE-associated sRNAs in lethal and viable paternal excess showed a pattern similar to that of bulk sRNAs (Supplemental Figure 4A,B). These observations suggest that the capacity to produce 24 nt sRNAs is relatively normal in viable paternal excess endosperm, presumably because of restored *RDR2* expression (Figure 4A,B). We also note that the proportion of 21 nt sRNAs increased in both lethal and viable paternal excess endosperm (Figure 4B, Supplemental Figure 4). While the pathways involved in the biogenesis of these 21 nt sRNAs remain unclear, they could represent the activity of post-transcriptional gene silencing (mRNA cleavage) pathways on transcripts that are mis-regulated in both lethal and viable paternal excess endosperm (Borges and Martienssen, 2015; Marí-Ordóñez et al., 2013)

A hallmark of the activity of the 24 nt sRNA pathway is CHH methylation. We performed whole genome bisulfite sequencing to profile the methylomes of balanced, lethal, and viable paternal excess endosperm (Supplemental Table 1). To identify regions with altered CHH methylation, we divided the genome into 300 bp windows with 200 bp overlaps and identified those windows with a 10% or greater CHH methylation difference between genotypes (i.e. 20% vs 30% methylation.) Windows with significant differences were merged into differentially methylated regions. CHH methylation was slightly higher in viable paternal excess endosperm relative to lethal paternal excess endosperm, but in both genotypes it was drastically reduced relative to balanced endosperm on both maternal and paternal alleles (Figure 4C, Supplemental Datasets 4 & 5). Regions with reduced CHH methylation significantly overlapped with regions with lowered sRNA levels in lethal paternal excess endosperm (Figure 4D).

### Paternal excess differentially affects the expression of maternally and paternally inherited alleles

Total gene expression levels are derived from the contribution of maternally and paternally inherited alleles. Based on the effect of *nrpd1* mutations on allele-specific expression in balanced endosperm, we previously proposed that 4N *nrpd1* might suppress paternal excess lethality by increasing transcriptional dosage from maternal alleles for many genes (Erdmann et al., 2017). To test this hypothesis, we evaluated allele-specific expression in balanced, lethal, and viable paternal excess endosperm for genes with at least 50 allele-specific reads in both balanced and paternal excess crosses (Figure 5) (Supplemental Dataset 9). Balanced endosperm has a 2:1 m:p ratio and paternal excess endosperm has a 2:2 ratio of maternal:paternal genomes. Mirroring the genomic ratios, the median paternal fraction for genes was 33.8% in balanced endosperm, 50.1% in lethal paternal excess endosperm, and 50.6% in viable paternal excess (Figure 5A). To assess potential allelic mis-regulation of individual genes, we normalized transcriptomic fractions to genomic fractions by calculating the deviation from paternal genomic contributions for each gene (% paternal-33% for balanced crosses and % paternal-50% for paternal excess). Contrary to our expectation, there was a small but detectable *decrease* in paternal contribution relative to genomic expectation for specific genes in lethal paternal excess endosperm (*two-sided D’Agostino’s K^2^ test, skew =−0.58*, *p< 2.2e-16*; Figure 5B, purple line, Supplemental Dataset 9). A similar bias was observed in viable paternal excess endosperm (*two-sided D’Agostino’s K^2^* test, *skew =−0.53*, *p <2.2e-16;* Figure 5B, yellow line, Supplemental Dataset 9). Thus, in viable relative to lethal paternal excess endosperm there were few changes in overall allelic bias (*two-sided D’Agostino’s K^2^* test, *skew = 0.0443*, *p = 0.05887;* Figure 5B, gray line, Supplemental Dataset 9).

**Figure 5:**
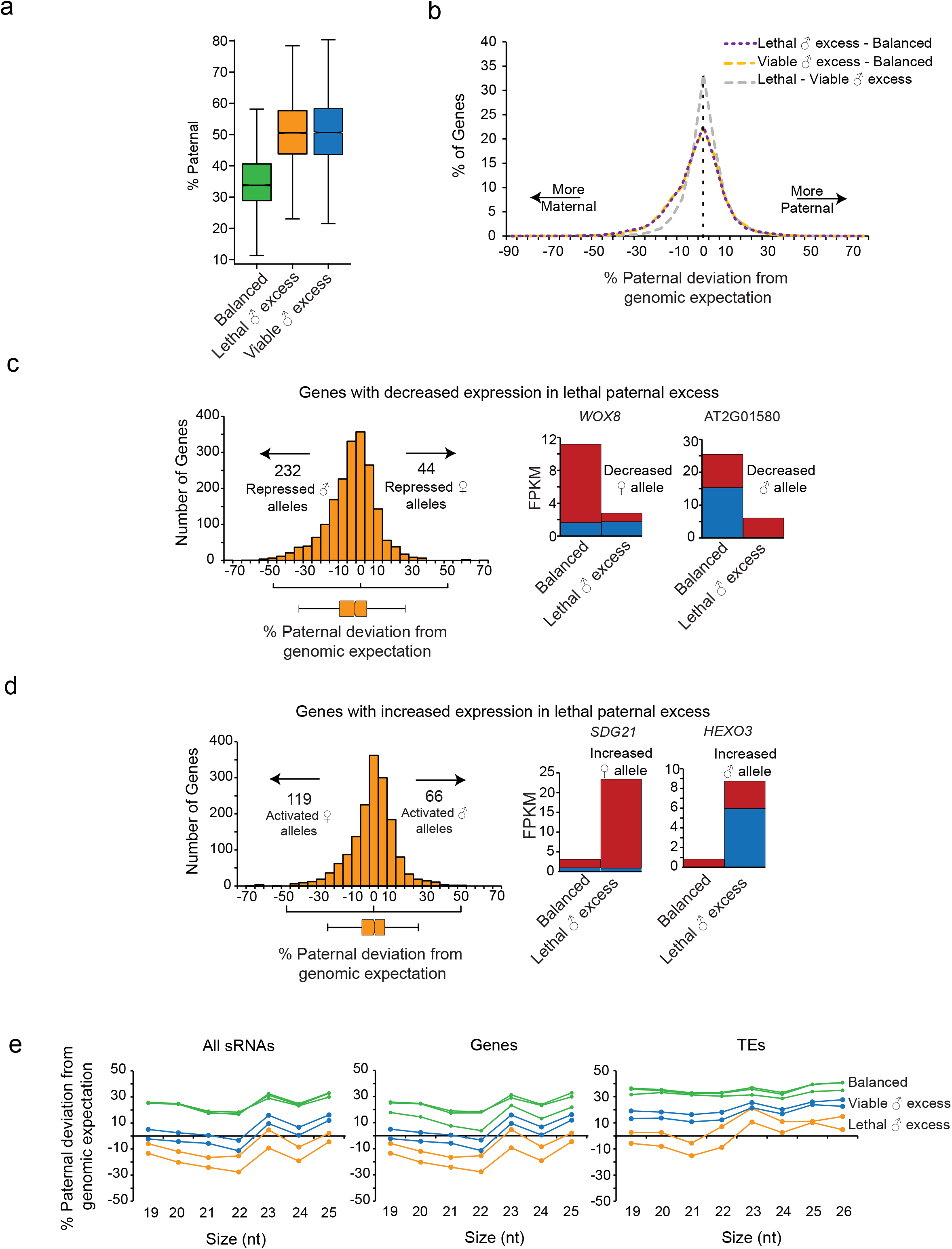
Allelic contributions to expression differences between balanced and lethal paternal excess endosperm. **a)** Paternal fraction in genic transcripts. Boxplot represents all genes with at least 50 allele specific reads in indicated genotype. Number of genes in balanced = 10,388, lethal ♂ excess =11,709, viable ♂ excess =11,430. For balanced endosperm, a total of five replicate libraries were analyzed (Erdmann et al., 2017). **b)** Paternal excess endosperm transcriptome is maternally skewed. Frequency distribution plot of % paternal deviation for each gene. Paternal deviation was calculated for genes with at least 50 allele specific reads in each pair of genotypes being compared. Number of genes in lethal ♂ excess - balanced = 9,550, viable ♂ excess - balanced = 9,639, viable ♂ excess - lethal ♂ excess = 10,919. **c)** Decreased paternal allele contribution at a larger proportion of genes with decreased expression in lethal paternal excess compared to balanced. N = 1910 genes. Impacts of allele-specific changes on gene expression at two examples, *WOX8* and *AT2G01580*, are shown. **d)** Increased maternal allele activation at a larger proportion of genes with increased expression in lethal paternal excess. N= 1608 genes. Impacts of allele specific changes on gene expression at two examples, *SDG21* and *HEXO3*, are shown. Genes analyzed for **c** and **d** were detected as being significantly different in gene expression by Cuffdiff. A gene with an allelic shift of at least 20% was considered to have a modified allelic balance. Boxplots represent median values for paternal deviation from genomic expectation. For gene-specific histograms, maternal allele is red and paternal allele is blue. **e)** sRNA populations are increasingly maternally biased in paternal excess endosperm. The paternal fraction of sRNA populations from viable paternal excess endosperm is intermediate between lethal paternal excess and balanced endosperm. Data from three replicates of balanced endosperm and two replicates each of lethal and viable paternal excess endosperm are plotted here. sRNA reads mapped to TAIR10 genome were first split based on size and then into Col and L*er* reads and reads using SNPs. % of paternal reads, and deviation from genomic expectation at each size was calculated.

To explore the contribution of allelic mis-regulation to genic mis-regulation, we examined shifts in paternal deviation from genomic expectation for genes with increased or decreased expression in lethal paternal excess endosperm compared to balanced endosperm (Figure 5C-D, Supplemental Figure 5A). Genes with decreased expression were more heavily influenced by loss of paternal allele contributions (*two-sided D’Agostino’s K^2^ test, skew for down-regulated genes =-0.4506, p-value= 1.243e-16;* Figure 5C). 12.1% of genes with sufficient allele-specific reads showed at least 20% decrease in paternal bias, indicating increased repression of paternal alleles, whereas 2.3% showed at least a 20% increase in paternal bias, indicating increased repression of maternal alleles (Figure 5C). Genes with increased expression in lethal paternal excess endosperm were more impacted by increases in maternal allele contributions (*skew for up-regulated genes=−0.52084, p-value= 6.61e-16*; Figure 5D). 7.4% showed at least a 20% decrease in paternal deviation, indicating increased maternal allele expression, and 4.1% showed at least a 20% increase in paternal bias, indicating increased paternal allele expression (Figure 5D).

Comparison of allelic contributions between viable and lethal endosperm revealed large differences at a very limited number of genes (Supplemental Figure 5B-D). Among genes with increased transcript abundance in viable compared to lethal paternal excess endosperm, increased paternal allele contributions were observed at 9 loci (Supplemental Figure 5B,C, Supplemental Dataset 9). Increased maternal allele contributions were observed at another 9 loci (Supplemental Figure 5B,C, Supplemental Dataset 9). Among genes with decreased abundance, only two genes showed strong evidence for a repression of paternal allele expression and two genes had evidence for repression of maternal allele expression (Supplemental Figure 5B,D).

We also tested if the increased expression of imprinted genes in paternal excess endosperm was related to a breakdown of imprinting (Supplemental Figure 6). In comparisons of balanced endosperm and lethal paternal excess, loss-of-imprinting was apparent among MEGs. Among 27 MEGs with increased expression in lethal paternal excess endosperm, paternal deviation increased for 8 genes, which became bi-allelically expressed (Supplemental Figure 6). However, strongly maternally biased expression was maintained at 11 MEGs (Supplemental Figure 6). Of 25 PEGs with increased total expression, 13 had reduced paternal bias (Supplemental Figure 6). However, most of these changes were small, with the exception of AT4G20800, which was bi-allelically expressed (Supplemental Figure 6). These observations indicate that loss-of-imprinting contributes to increases in imprinted gene expression (Figure 3) at only a subset of imprinted genes.

The allele-specific changes in gene expression (Figure 5A-D) led us to test if sRNAs change in an allele-specific manner. The sRNA pool in balanced endosperm is overall paternally biased (Erdmann et al., 2017) (Figure 5E). For most sRNA size classes, lethal paternal excess endosperm was more maternally-biased than expected from the ratio of maternal and paternal genomes (Figure 5E). For example, in lethal paternal excess, 24 nt sRNAs were 13.8% more maternal than expected given the 2:2 ratio of maternal to paternal genomes. This indicates, counterintuitively, that extra paternal genomes are associated with decreased small RNAs from paternal genomes (or increased sRNAs from maternal genomes). sRNAs in viable paternal excess endosperm are slightly more paternal than lethal paternal excess endosperm, although to a lesser extent than is observed in balanced endosperm (Figure 5E), consistent with increased functionality of the RdDM pathway in viable paternal excess endosperm (Figure 4A).

In summary, from these analyses we conclude that increasing paternal genome copy number can impact the expression of only maternal or only paternal alleles for a subset of loci. Thus, maternally and paternally inherited alleles contribute unequally to genome-wide transcriptome mis-regulation. However, overall, we did not find evidence for our hypothesis that a shift toward maternal allele expression is associated with repression of paternal excess seed abortion (Erdmann et al., 2017), as *both* lethal and viable paternal excess endosperm have a higher fraction of maternal transcripts than expected from parental genomic ratios.

## DISCUSSION

### Impact on models for interploidy seed abortion

We have shown that an extra copy of the paternal genome induces regulatory changes at both maternal and paternal alleles and drives massive changes in gene expression in the endosperm, varying little between viable and inviable seeds (Figures 2-5). Our data allow evaluation of a number of proposals regarding the transcriptional changes at genes, TEs, or sRNAs that could result in interploid seed lethality. Inspired by models of transposon-induced dysgenesis in *Drosophila* leading to atrophied ovaries (reviewed in Kelleher, 2016), it was speculated that an imbalance between sRNAs deposited by a diploid mother and the TE load in a tetraploid father could lead to TE expression, which would then trigger seed abortion (Castillo and Moyle, 2012; Martienssen, 2010). Our finding that TEs are similarly mis-regulated in lethal and viable paternal excess endosperm (Figure 3) leads us to conclude that TE expression levels are unlikely to be a determinant of interploidy seed viability. While it is perhaps surprising that viable seeds have high TE transcript abundance, it is not unprecedented. Arabidopsis mutants like *met1*, which have high levels of TE expression (Oberlin et al., 2017; Zilberman et al., 2007), also produce viable seed (Xiao et al., 2006).

Transcription of imprinted genes is potentially a strong candidate for the critical gene dosage difference separating lethality and viability (Haig and Westboy, 1991; Lafon-Placette and Köhler, 2015; Wolff et al., 2015). But, like TEs, the similar expression levels of imprinted genes between lethal and viable interploid seeds (Figure 3) suggest their differential expression is not determinant for viability. We did identify 614 differentially expressed genes between lethal and viable interploidy seeds (Figure 2B,C, Supplemental Dataset 1). These 614 differentially expressed genes include several genes encoding proteins involved in phytohormone signaling and developmental regulators – all potentially good candidates for determining seed viability (Supplemental Dataset 1). It is possible that the expression of multiple loci of modest effect or partial renormalization of expression patterns of multiple loci constitute the dosage threshold between lethal and viable seeds. Based on the present data, we are unable to distinguish if any of the differentially expressed genes or partially corrected genes are causal for the effect on seed viability, or whether they represent a consequence of endosperm development associated with seed viability.

Several distinct hypotheses revolving around sRNAs have been postulated to explain how loss of *NRPD1* in tetraploid fathers, but not diploid mothers, represses seed abortion in paternal excess crosses (Erdmann et al., 2017; Martinez et al., 2018). The easiRNA model for interploid seed abortion is based on two key conclusions from the recent data of Martinez et al (Martinez et al., 2018). First, a non-canonical RNA Pol IV-RDR6 pathway functions in pollen to make easiRNAs, which scale with paternal genome dosage. Second, the transfer of increased easiRNAs into the endosperm from diploid pollen inhibits RdDM; it is then the absence of these easiRNAs in *nrpd1* mutant pollen that permits the RdDM pathway to function in paternal excess endosperm. Our data do not support this model. We failed to obtain a significant increase in viable paternal excess seeds with 4N *rdr6-15* pollen (Figure 1A, Supplemental Figure 1B), suggesting that any sRNAs exclusively dependent on *RDR6* are not linked to paternal excess seed abortion. Consistent with this result, paternally produced siRNA854, which is dependent on *DCL2/4* and *RDR6*, is required for viability of paternal excess seeds (McCue et al., 2012; Wang et al., 2018). By contrast, we found that multiple members of the canonical RdDM pathway restore paternal excess seed viability (Figure 1, Supplemental Figure 1A,C). Unlike the reported relatively low levels of viable seed produced by mutations in non-RdDM genes such as *RDR6, DCL2, DCL4*, *AGO2* and *miRNA845* (approximately 20-30% of the seeds are viable) (Borges et al., 2018; Martinez et al., 2018), loss of *NRPE1* and *DRM2* phenocopy *NRPD1* in the large magnitude of repression of seed abortion (approximately 80% of the seeds are viable) (Figure 1, Supplemental Figure 1). The cause of the discrepancy among the genetic results between our studies remains unclear, but may be linked to the use of *osd1* mutation to create conditions of paternal excess. Similar discrepancies have been observed elsewhere; *osd1 pkr2* mutants did not repress paternal excess seed abortion whereas 4N *pkr2* showed reduced seed abortion (Huang et al., 2017; Wolff et al., 2015). Finally, the easiRNA model also suggests that the RdDM pathway’s recovery in the viable paternal excess endosperm is essential for seed viability. However, loss of both paternal and maternal copies of *NRPD1, RDR2, DCL3, NRPE1*, and *DRM2* in paternal excess crosses did not abolish the seed viability obtained upon loss of the paternal copy alone (Figure 1, Supplemental Figure 1) (Erdmann et al., 2017; Martinez et al., 2018). This indicates that the activity of a genetically complete canonical RdDM pathway in the endosperm is not necessary for suppression of paternal excess seed abortion.

Another model posits that RNA Pol IV sRNA levels are determined by maternal dosage and that these small RNAs normally repress *AGL*s and trigger endosperm cellularization (Lu et al., 2012). This model argues that in paternal excess endosperm siRNA levels are disproportionate to target transcript levels of *AGL* genes, which in turn leads to prolonged endosperm proliferation and cellularization. This model makes a key testable prediction – the requirement for a functional RdDM pathway in the endosperm. However, we find that RdDM in endosperm is not essential for seed viability in conditions of paternal excess (Figure 1, Supplemental Figure 1).

We previously showed that maintaining a 2:1 ratio of maternal to paternal allele transcripts in balanced endosperm was dependent on *NRPD1*; in its absence the expression of many genes shifted more maternal (Erdmann et al., 2017). Thus, we suggested that the loss of *NRPD1* repressed paternal excess seed abortion by raising the maternal fraction of the transcriptome and balancing excess paternal gene dosage (Erdmann et al., 2017). A central tenet of our model was that the reduction in the RNA Pol IV sRNA pathway in endosperm (caused by loss of paternal *NRPD1*) was essential for seed viability. However, we find that the RdDM pathway is in fact *more* functional in viable paternal excess endosperm than in lethal paternal excess endosperm (Figure 4A-B, Figure 5E, Supplemental Figure 4). Additionally, the transcriptome in lethal paternal excess is already maternally biased relative to genomic expectation. This maternal bias is not increased in viable paternal excess endosperm, as would be predicted under our previous model (Figure 4).

What, then, might be the mechanism by which suppression of seed lethality occurs? Although our current data cannot address whether mutations in all canonical RdDM genes suppress paternal excess seed lethality in the same manner, given what is known about the RdDM pathway it is likely that differences in DNA methylation, in either in the paternal sporophyte or the male gametophyte (or both), play a key role. It has been shown in diploid plants that loss of *NRPD1* or *DRM2* in the father can impact gene expression in the endosperm of balanced crosses (Erdmann et al., 2017; Vu et al., 2013) and that *DRM2* establishes methylation patterns at a subset of genomic sites in the male germline (Walker et al., 2018). Some studies suggest that tetraploidization itself induces DNA methylation changes (Baubec et al., 2010; Mittelsten Scheid et al., 2003; Yu et al., 2010; Zhang et al., 2015). In one scenario, DNA methylation changes in tetraploid pollen could ultimately direct changes in gene expression in the endosperm, which would not occur in endosperm fathered by RdDM mutant tetraploids. An alternative possibility is that loss of RdDM in tetraploids could lead to an altered methylation state not found in either wild-type diploid or tetraploid males. This ectopic paternal methylation state could then dictate a gene expression pattern that represses seed abortion.

### Evidence for buffering mechanisms in endosperm

Our results suggest that transcriptional buffering, operational in both lethal and viable paternal excess endosperm, is one feature that could contribute to the ability of the seed to withstand the addition of an entire extra paternal genome. In transcriptional buffering, expression per copy of a gene can be increased or decreased based on the number of copies of the gene relative to that of the rest of the genome (Birchler and Veitia, 2012; Zhang et al., 2010). We observed the signatures of buffering in paternal excess transcriptomes (Figure 5). Decreased paternal transcript levels contributed predominantly to genes with decreased expression in paternal excess, whereas increases in the levels of transcripts from maternal alleles contributed predominantly to genes with increased expression (Figure 7C-D). The outcome is that some parts of the transcriptome are more maternal than expected from parental genomic dosage, constituting a push back against excess paternal genomes. While the mechanistic basis of this buffering system remains unclear, it could be linked at least in part to the reduction in the 24 nt small RNA pathway in endosperm (Figure 4). RNA Pol IV function in balanced endosperm has been previously shown to be required for repression of the maternal transcriptome (Erdmann et al., 2017). The increase in the maternal fraction of the transcriptome for specific genes in paternal excess endosperm (Figure 5B) is therefore consistent with the reduced expression of genes encoding subunits of RNA Pol IV and other downstream members of the RdDM pathway (Figure 4A). It is important to note that the increase in maternal fraction can be observed in both lethal and viable paternal excess endosperm and is thus unlinked to the repression of seed abortion brought about by paternal loss of RdDM. Other buffering mechanisms might also play important roles and could act through post-transcriptional pathways (Wang et al., 2018), protein degradation pathways, or via signaling pathways that have been shown to influence paternal excess crosses (Batista et al., 2019; Doughty et al., 2014). One striking observation is that even wild-type paternal excess crosses produce about 20% viable seed (Figure 1, Supplemental Figure 1), suggesting that seed lethality and viability exist on a continuum. Our data further suggest that the slim margin that separates seed life and death is likely determined by modest changes in the endosperm, which are ultimately instructed by paternal imprints laid down by RdDM.

## Methods

### Arabidopsis strains, tetraploid production, and tissue collection

Mutants used in this study were: *nrpd1a-4*, *rdr2-1*, *dcl3-1*, *nrpd1b-11*, *drm2-2*, *rdr6-15* (CS879578), *rdr2-1 rdr6-15* (CS66111), and *cmt3-11t* (CS16392) and were obtained from ABRC. Tetraploids were generated by applying 0.25% colchicine in 0.2% Silwet to the apices of 2-3 week old diploid plants. Progeny from treated plants were screened by flow cytometry to identify tetraploids. For crosses, wild-type diploid Col-0 or L*er* buds were emasculated and pollination was carried out two days later with diploid or tetraploid pollen in Col-0 strain background. To assess seed abortion, siliques were harvested after drying and seeds examined under a dissecting microscope. To assess germination rates, seeds were sterilized in 2% PPM (Plant Cell Technologies) for three days at 4°C and then plated on 0.5X MS/Phytagar media. Endosperm from approximately one hundred seeds at 7 DAP from at least three siliques were dissected away from embryo and seed coat and pooled for each replicate as previously described (Gehring et al., 2009; 2011). Replicates were collected from a different sets of crosses with different individuals as parents. For lethal paternal excess seeds, endosperm was collected only from seeds with arrested embryonic growth. For viable paternal excess seeds, endosperm was collected only from seeds where embryonic growth was progressing normally.

### mRNA-Seq library preparation and transcriptome analyses

Long RNA was isolated as previously described (Erdmann et al, 2017) and mRNA-seq libraries were constructed at the Whitehead Institute Genome Technology Core using the SMARTerUltra-lowPOLYA-V4 kit. Libraries were sequenced on a 40 base, single read cycle. Sequence data were filtered for quality with “ *trim_galore -q 25 --phred64 -- fastqc --length 20 --stringency 5*”. Filtered reads were aligned to the TAIR10 genome using “*tophat -i 30 -I 3000 --solexa1.3-quals -p 5 -g 5 --segment-mismatches 1 -- segment-length 18 --b2-very-sensitive*” (Kim et al., 2013). Tophat version 2.1.1 with Bowtie version 2.3.4 was used. We used Cuffdiff (version 2.2.1) with default settings and the ARAPORT11 genome annotation to calculate changes in gene expression and their statistical significance. Genes with q-value <0.05 were considered to be significantly different. We used a custom script (assign_to_allele.py; https://github.com/clp90/imprinting_analysis/tree/master/helper_scripts) and SNPs between Col-0 and L*er* to identify allele specific reads. Allele specific reads per gene were assessed using “*htseq-count -s no –m union*” and the ARAPORT11 annotation. To assess TE transcript levels, reads were aligned to the REPBASE consensus sequence (Jurka et al., 2005) using “*bowtie -v 2 -m 3 --best --strata -p 5 --phred64”* and reads mapping to each TE family were summed. *META1, ATHILA6A* and *ATGP8* were precluded from further analyses because of mapping artifacts. Differential abundance of TE mapping reads were calculated using Fisher’s exact test (p<0.05) option with multiple-test correction in DEGseq (Wang et al., 2010).

### Tissue enrichment test

Expression per gene was measured using HT-Seq count and analyses were performed using the seed tissue enrichment test (Schon and Nodine, 2017). The time point was set to bent cotyledon.

### Small RNA-Seq library preparation and analyses

RNA was isolated from manually dissected endosperm using the RNAqueous micro kit (Ambion). Small RNA was obtained as previously described (Erdmann et al, 2017). Libraries were built using the NEXTflex sRNA-seq kit v3 (Bioo Scientific). Final library amplification was carried out for 24 cycles; resultant libraries were size selected (135- 160 bp) with a pippin prep. 40 base single read sequencing was carried out an Illumina HiSeq 2500. Sequencing reads were trimmed for quality with “*fastq_quality_trimmer -v - t 20 -l 25*”. Reads were further filtered and adapter containing reads were retained using “*cutadapt -a TGGAATTCTCGGGTGCCAAGG --trimmed-only --quality-base 64 -m 24 - M 40 --max-n 0.5”*. The reads from libraries produced by the NEXTflex library prep kit include four random nucleotides at both 5’ and 3’ ends of the reads. Taking advantage of these tags, we were able to remove PCR duplicates using “*prinseq-lite –fastq <infile> -out_format 3 –out_good <outfile> -derep 1”*. Reads were aligned to TAIR10 with “*bowtie -v 1 --best -5 4 -3 4 --sam --phred64-quals or --phred33-quals*”. We used a custom script (assign_to_allele.py; https://github.com/clp90/imprinting_analysis/tree/master/helper_scripts) to identify allele-specific reads. Regions with differential sRNA levels were identified by counting reads in 300 bp windows with 200 bp overlaps. DESeq2 (Love et al., 2014) was used to identify windows with differential abundance of sRNAs (adjusted *p-value* less than or equal to 0.05 and log_2_ fold change =+/-1). Overlapping windows with significant increases or decreases in sRNA were merged to obtain regions with differences in sRNA. To identify genes with differences in 21 and 24 nt sRNAs, mapped reads were separated based on size. 21 and 24 nt sRNAs mapping to genes were counted using the ARAPORT11 annotation and “*htseq-count -s no –m union*”. To identify TE insertions with differences in 21 and 24 nt sRNAs, we first identified TE insertions that did not overlap with genes and used “*coverageBed -counts*” command from Bedtools suite (Quinlan and Hall, 2010) to count reads mapping to TE insertions. We identified genes and TEs with differential abundance in 21 and 24 nt sRNA using DESeq2.

### Methylome library preparation and analyses

Genomic DNA was isolated from manually dissected endosperm using the QiaAMP DNA microkit (QIAGEN 56304); dissected tissue was incubated in ATL buffer and proteinase K at 56°C overnight on a shaking incubator. 80-100 ng of DNA obtained from these protocols was subjected to bisulfite treatment using Methylcode Bisulfite Conversion kit (Invitrogen). Bisulfite libraries were constructed from these materials using Illumina’s Truseq DNA methylation kit. Bisulfite libraries were sequenced on Illumina’s HISeq 2500 in a paired end configuration. Reads were filtered for quality with “*trim_galore --phred64 --fastqc --stringency 5 --length 15 --paired --clip_R1 2 --clip_R2 2*”. The reads were then aligned to TAIR10 using *“bismark -N 1 -L 20”*. Duplicate reads were removed using Bismark version 0.19. Bismark methylation extractor and custom scripts previously described (Pignatta et al., 2014, 2015) were used to obtain per base methylation. Briefly, differences in methylation were calculated genome-wide for 300 bp sliding windows with 200 bp overlaps. To be included, windows had at least 3 overlapping cytosines in both genotypes with a read depth of at least 6 reads per cytosine. Windows called as significantly different were at least 10% different between tested genotypes for CHH methylation, 20% for CHG methylation and 30% for CG methylation. Significance of difference was calculated by F.E.T with a Bonferroni-Hochberg correction (p<0.01). Overlapping windows with significant differences in DNA methylation were merged to obtained differentially methylated regions.

### Data Access

Whole genome bisulfite sequencing data, small RNA sequencing and mRNA sequencing data are deposited in NCBI GEO under accession GSE126932.

## Acknowledgements

We thank Dr. Daniela Pignatta for deriving the 4N Col-0 line. This research was supported by NSF MCB CAREER Award 1453459 to M.G.

## Author Contributions

P.R.V.S and M.G. conceived of and designed the study, P.R.V.S. performed experiments and analyzed data, and P.R.V.S. and M.G. wrote the manuscript.

## Supplemental Data Files

**Supplemental Dataset 1:** Cuffdiff output comparing gene expression in balanced, lethal and viable paternal excess endosperm.

**Supplemental Dataset 2:** Gene ontology analysis of genes that are differentially expressed between balanced, lethal and viable paternal excess endosperm.

**Supplemental Dataset 3:** DEGseq output comparing transposon transcript levels between balanced, lethal and viable paternal excess endosperm.

**Supplemental Dataset 4:** Differentially methylated regions in the CHH context in comparisons between balanced, lethal and viable paternal excess endosperm.

**Supplemental Dataset 5:** Differentially methylated region in the CHG context in comparisons between balanced, lethal and viable paternal excess endosperm.

**Supplemental Dataset 6:** Differentially methylated region in the CG context in comparisons between balanced, lethal and viable paternal excess endosperm.

**Supplemental Dataset 7:** DESeq output for differential abundance of 21 and 24 nt genic sRNA between balanced, lethal and viable paternal excess endosperm.

**Supplemental Dataset 8:** DESeq output for differential abundance of 21 and 24 nt TE sRNA between balanced, lethal and viable paternal excess endosperm.

**Supplemental Dataset 9:** Allele-specific mRNA read counts and differences in allelic contributions between balanced, lethal and viable paternal excess endosperm.

**Supplemental Figure 1:**
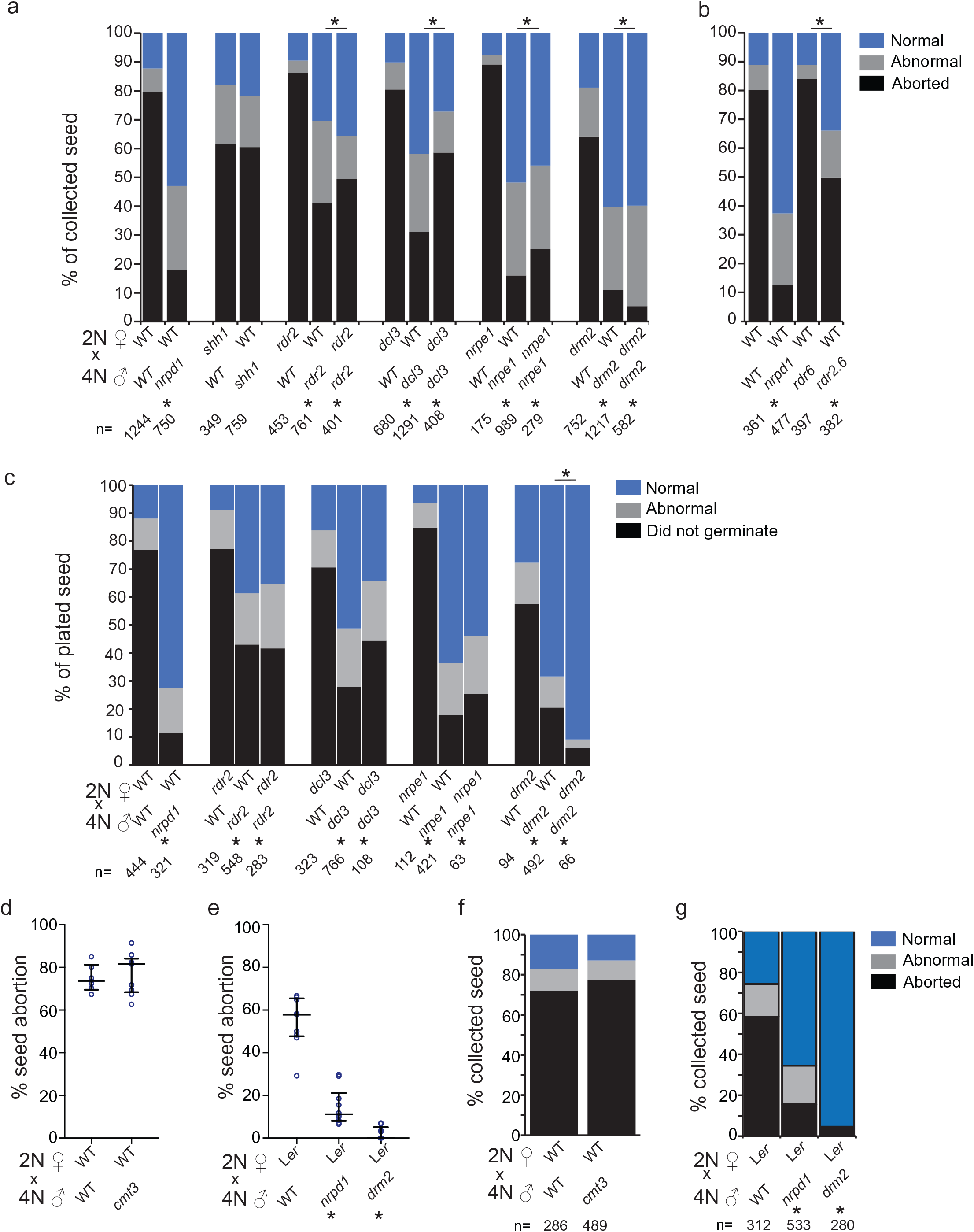
Loss of paternal RdDM genes but not *RDR6* and *CMT3* represses seed abortion in paternal excess crosses. **a,b,f,g)** Histograms depicting percentage of seed collected that was either aborted, abnormal or normal. **c)** Histograms depicting the germination status of collected seed. Seeds grouped under “normal” produced both true leaves and roots. Abnormally germinating seeds germinated but failed to produce a radicle or had abnormal cotyledons. Seeds that failed to produce either a radicle or a hypoctyl are grouped under “did not germinate”. **d)** Loss of CMT3 does not repress paternal excess lethality. Wilcox test did not show statistically significant differences between crosses in which the tetraploid father is either wild-type or *cmt3*. **e)** Interploidy crosses with L*er* mothers are similar to crosses with Col-0 mothers. Wilcox test showed statistically significant differences between crosses. For **d** and **e**, circles represent a single cross and includes multiple siliques from a single inflorescence. Bars represent median and interquartile range. * at bottom represents statistically significant difference in seed abortion or germination (p<0.05) in comparisons between indicated cross and cross between wild-type diploid mothers and wild-type tetraploid fathers. Statistical significance calculated by Chi-Squared test. N represents number of seeds/seedlings counted in each cross. WT = Col-0. This supplement supports Figure 1.

**Supplemental Figure 2:**
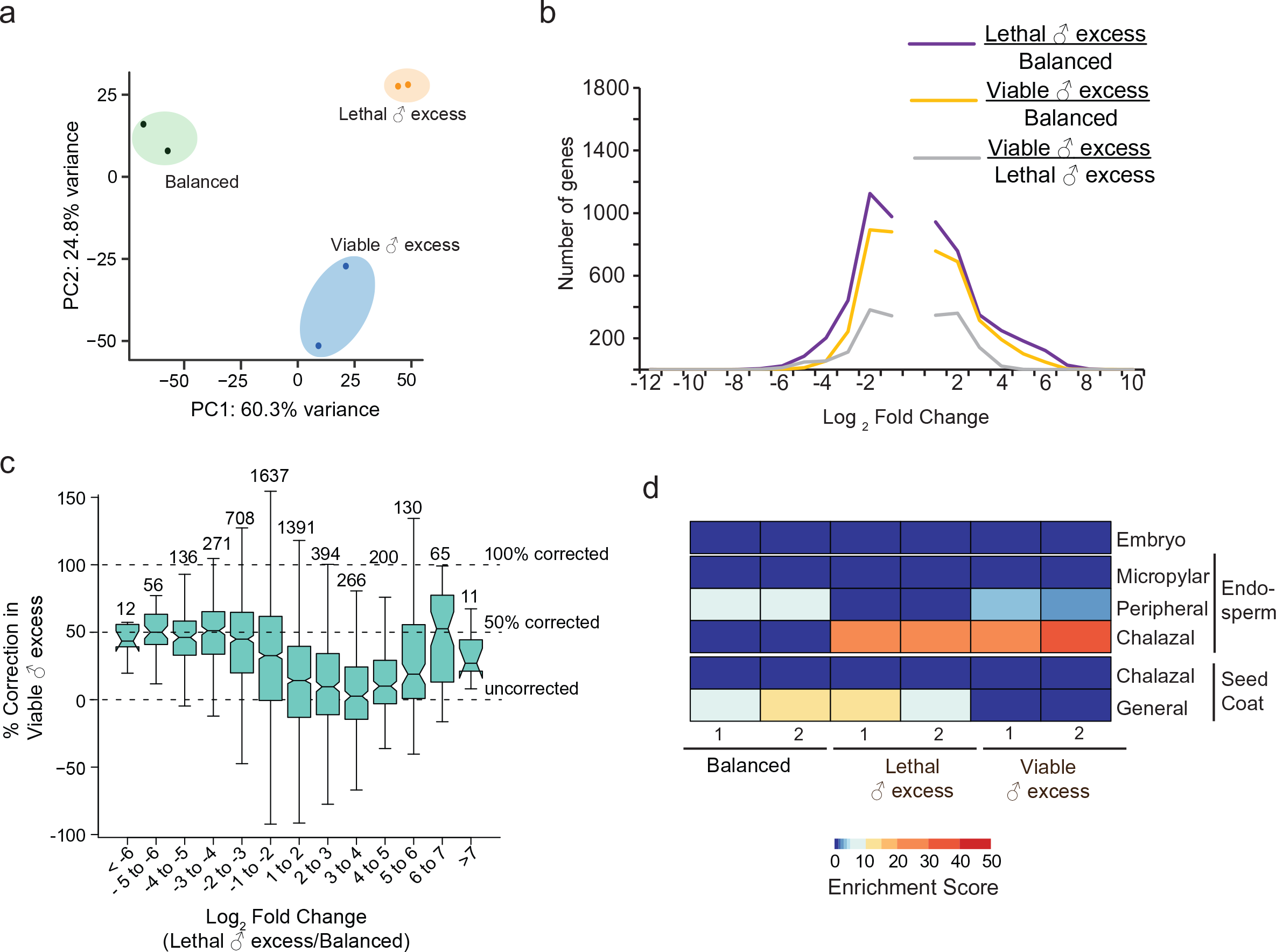
Analysis of gene expression differences in ♂ excess endosperm derived using the *osd1* mutation (data from Martinez et al., 2018). a) PCA plot shows that the transcriptomes of L*er* x Col *osd1* (lethal paternal excess) and L*er* x Col *osd1 nrpd1* (viable paternal excess) are more similar to each other than to L*er* x Col (balanced). b) Frequency distribution of all gene expression changes between balanced, lethal ♂ excess and viable ♂ excess. Fold change was calculated using Cuffdiff. All genes whose output FC was greater than 1 or less than -1 shown. c) Correction value was calculated for genes that are statistically significantly different in comparisons of balanced and lethal ♂ excess endosperm. The value, calculated as % Correction= 100-((log_2_ (viable/balanced))/(log_2_ (lethal/balanced))*100). Value of 100% indicates that a gene misregulated in lethal ♂ excess is not misregulated in viable ♂ excess. 0% represents similar misregulation in both lethal and viable ♂ excess. Fold change values and significance were calculated using Cuffdiff. Boxplot here is a Tukey plot. d) Both L*er* x Col *osd1* and L*er* x Col *osd1 nrpd1* endosperm are enriched for chalazal endosperm marker gene expression. Analysis was performed using the tissue enrichment test (Schon and Nodine, 2017) on gene expression measured by the HT-Seq count command. Tissue enrichment per biological replicate shown. This supplement supports Figure 2.

**Supplemental Figure 3:**
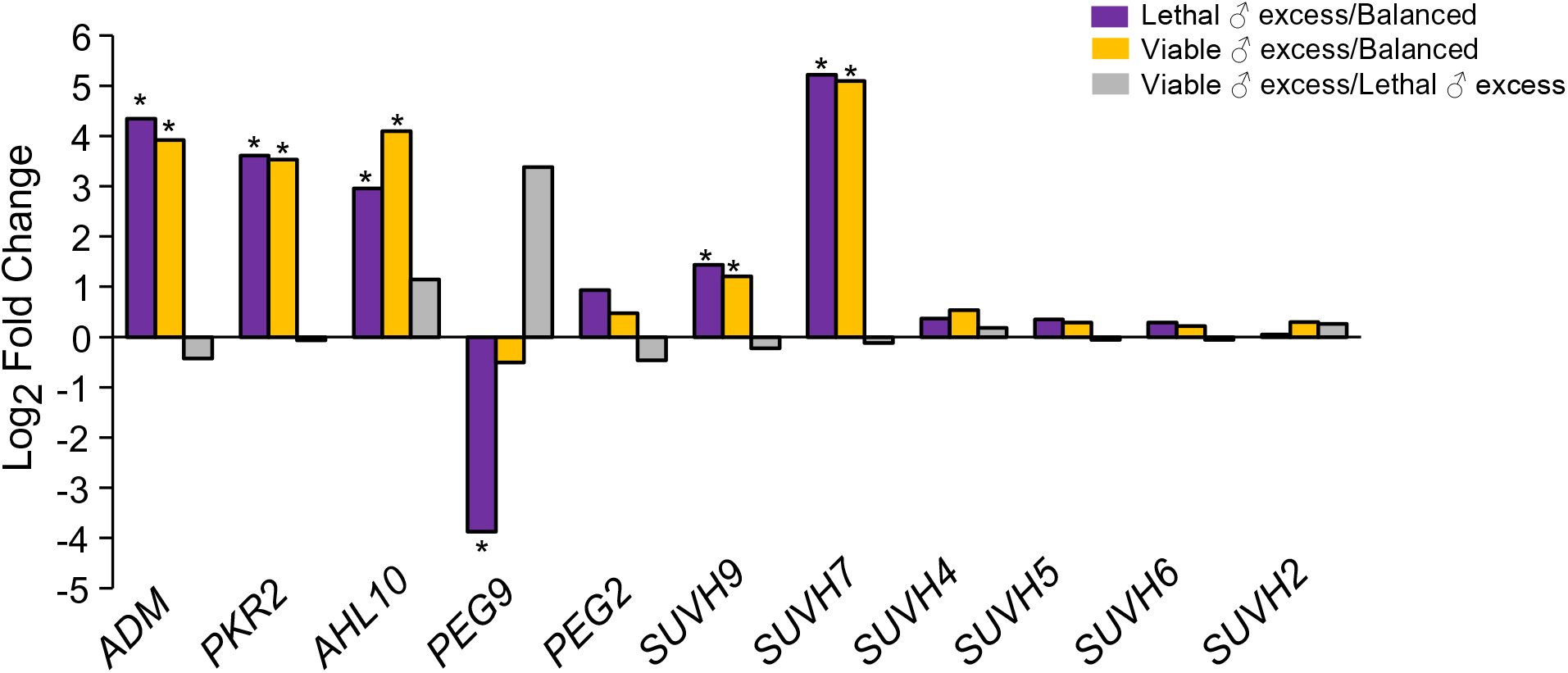
Suppression of paternal excess lethality by loss of *NRPD1* does not require normalization of genes implicated in interploid seed abortion. Expression level of genes shown to repress seed abortion in *osd1*-based paternal excess crosses is not repressed by loss of *NRPD1*. Fold change and statistical significance (represented by *) calculated by Cuffdiff. This supplement supports Figure 3.

**Supplemental Figure 4.**
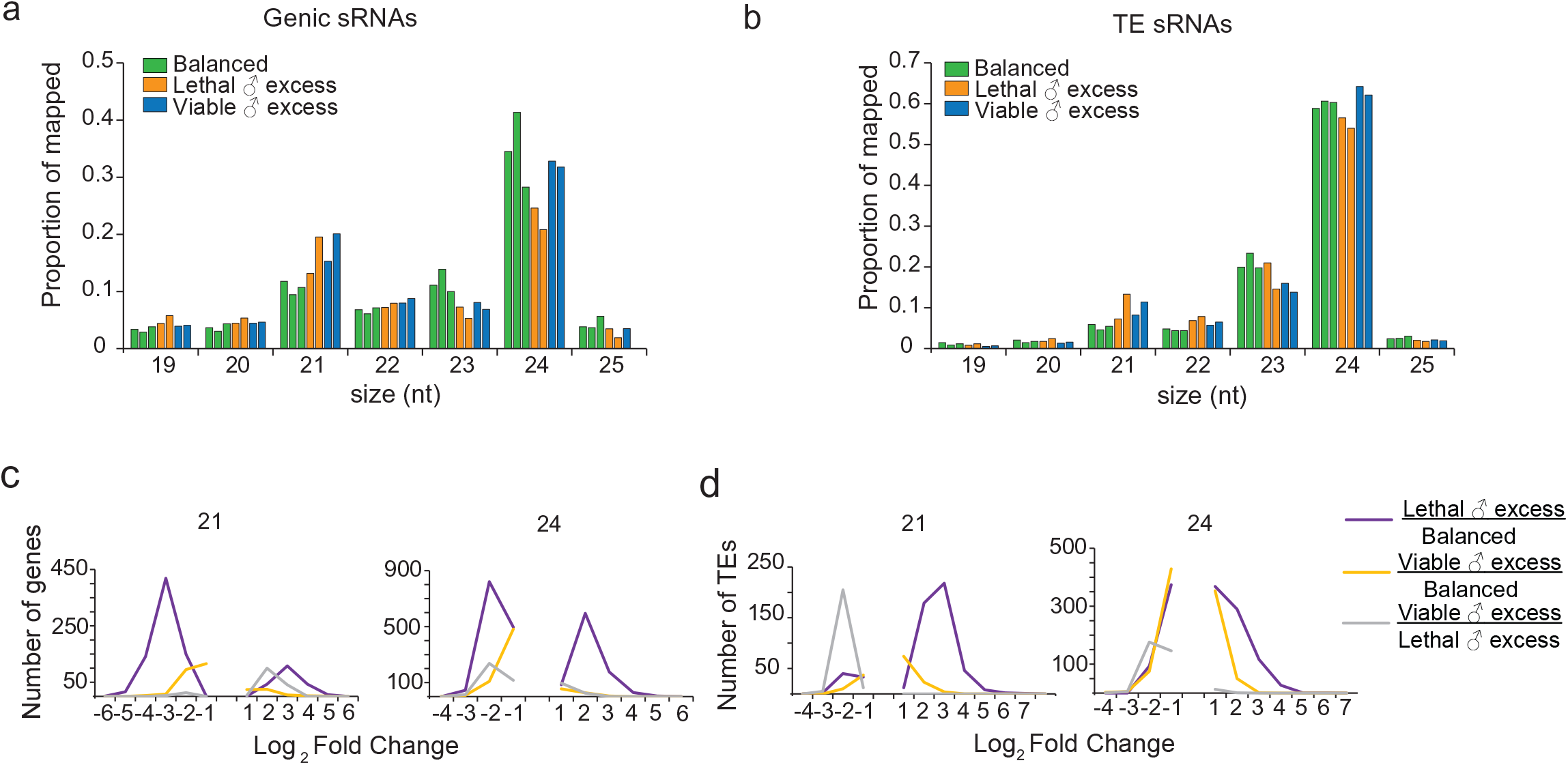
Small RNA in balanced, lethal and viable paternal excess endosperm. **a)** Genic sRNAs are reduced in lethal paternal excess. Size profiles of sRNA reads overlapping genes for three replicates of balanced endosperm, two replicates of lethal paternal excess and two replicates of viable paternal excess endosperm. Sum of all reads overlapping genes was used as denominator to determine proportion of small RNA at each size. **b)** TE insertion sRNAs are reduced in lethal paternal excess. Analysis as in a). Frequency distribution of significant changes in sRNA abundance at **c)** genes and **d)** transposable element insertions. Fold-change and significance calculated using DESeq2. Read counts for TAIR10 TEs generated using bedtools. This supplement supports Figure 4.

**Supplemental Figure 5:**
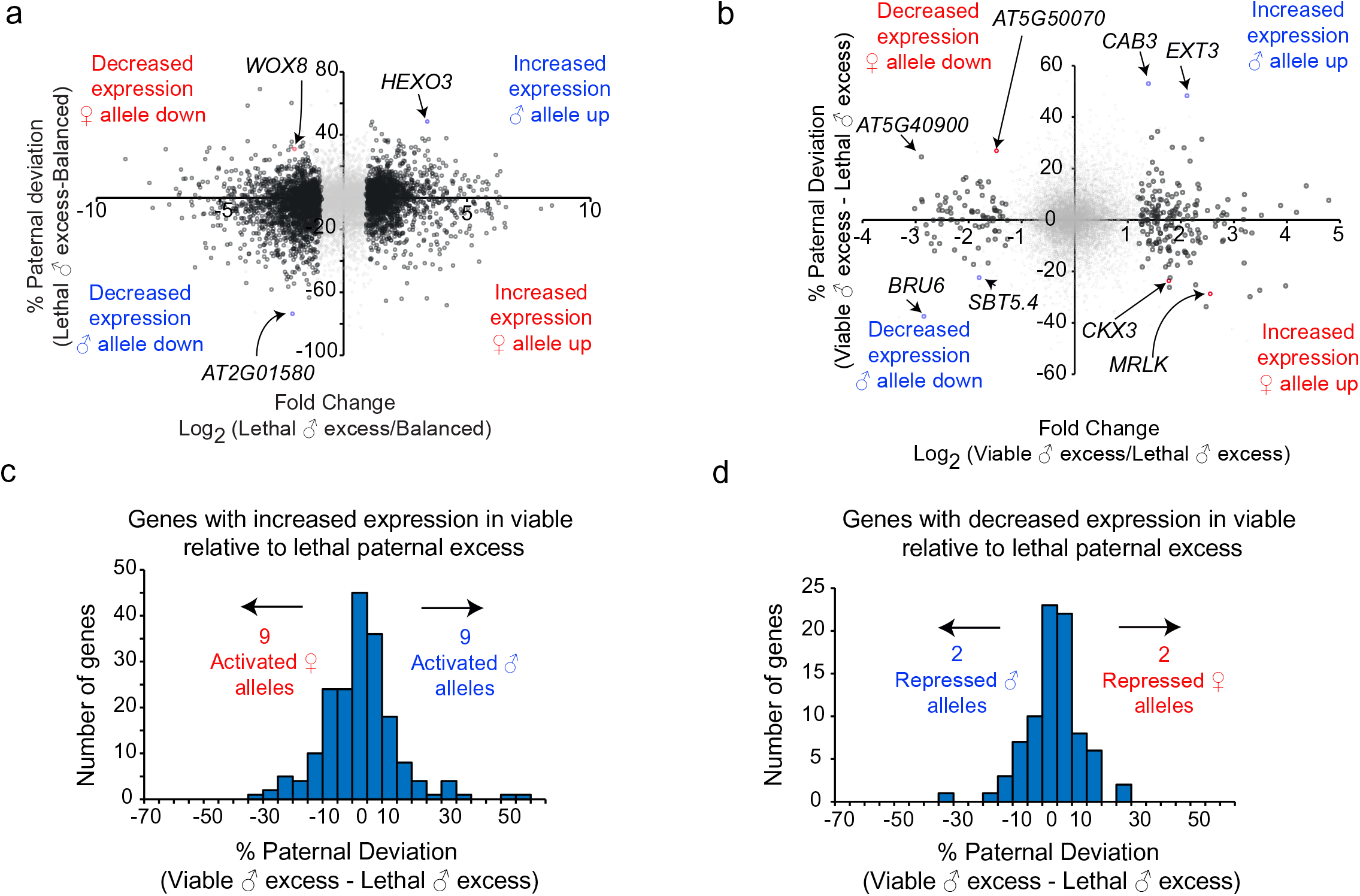
Allelic contributions to gene expression differences. **a)** Contribution of allelic mis-regulation to differential gene expression in lethal ♂ excess. Scatter plot of relationship between change in paternal deviation and change in gene expression between balanced and lethal paternal excess. **b)** Contribution of allelic mis-regulation to differential gene expression between lethal and viable ♂ excess. Scatter plot of relationship between change in paternal deviation and change in gene expression between balanced and lethal paternal excess. In **a** and **b**, black circles represent genes called as being significantly different by Cuffdiff; gray cricles represent genes that are not significantly different. **c)** Allele-specific changes in genes upregulated in viable relative to lethal paternal excess (N=189). **d)** Allele specific changes among genes downregulated in viable paternal excess relative to lethal paternal excess. (N=83) In **a-d**, genes with more than 50 allele specific reads were considered for analysis. In **c** and **d**, genes with at least a 20% shift in allelic balanced between viable and lethal ♂ excess were considered to have a modified allelic contribution. This supplement supports Figure 5.

**Supplemental Figure 6:**
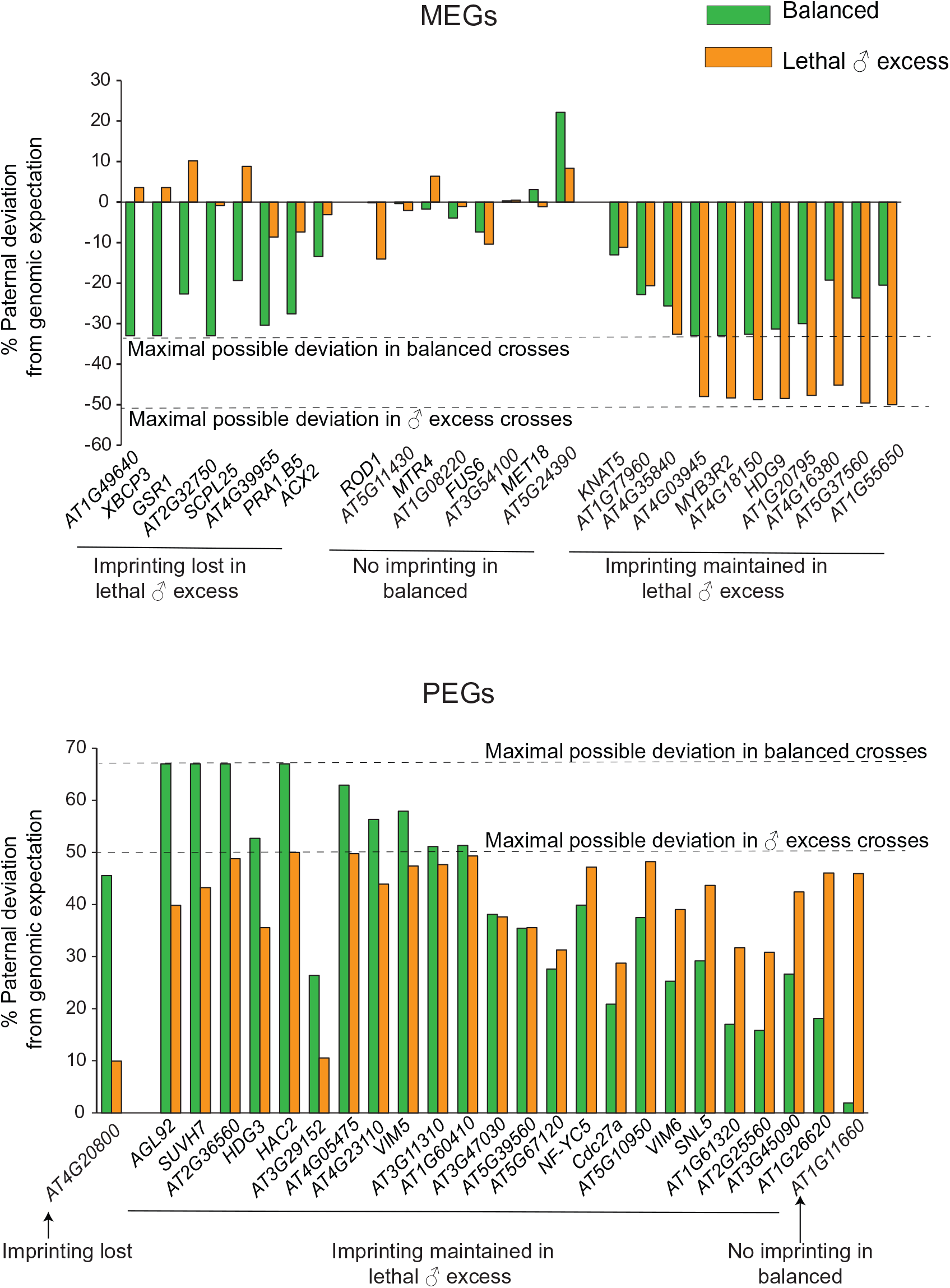
Allelic bias of some imprinted genes is lost in lethal paternal excess. Changes in paternal deviation for differentially expressed MEGs and PEGs. Maximal deviation for a MEG in balanced endosperm and paternal excess endosperm is -33 and -50 respectively. Maximal deviation for PEGs in balanced endosperm and paternal excess endosperm is 67 and 50 respectively. Our data did not show imprinting in balanced endosperm at a small subset of imprinted loci previously shown to be imprinted in balanced endosperm. This supplement supports Figure 5.

**Supplemental Table 1:**
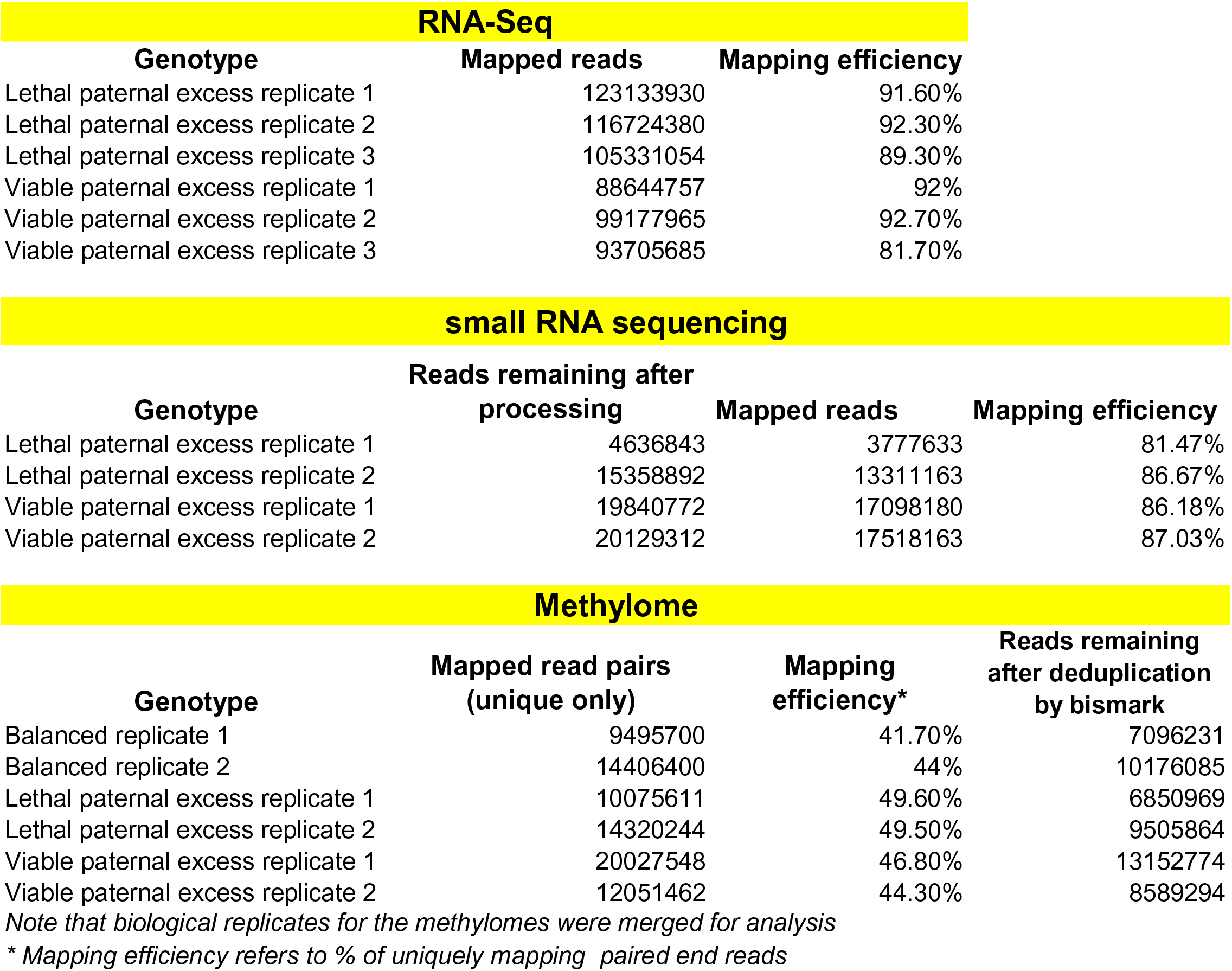
Details of sequencing libraires.

## References

Bao, W., Kojima, K.K., and Kohany, O. (2015). Repbase Update, a database of repetitive elements in eukaryotic genomes. Mob. DNA 6, 11.

Batista, R.A., Figueiredo, D.D., Santos-González, J., and Köhler, C. (2019). Auxin regulates endosperm cellularization in Arabidopsis. Genes Dev. 10.1101/gad.316554.118

Baubec, T., Dinh, H.Q., Pecinka, A., Rakic, B., Rozhon, W., Wohlrab, B., von Haeseler, A., and Scheid, O.M. (2010). Cooperation of Multiple Chromatin Modifications Can Generate Unanticipated Stability of Epigenetic States in Arabidopsis. Plant Cell 22, 34–47.

Birchler, J.A. (2014). Interploidy hybridization barrier of endosperm as a dosage interaction. Front. Plant Sci. 5.

Birchler, J.A., and Veitia, R.A. (2012). Gene balance hypothesis: Connecting issues of dosage sensitivity across biological disciplines. Proc. Natl. Acad. Sci. 109, 14746–14753.

Borges, F., and Martienssen, R.A. (2015). The expanding world of small RNAs in plants. Nat. Rev. Mol. Cell Biol. 16, 727–741.

Borges, F., Parent, J.-S., Ex, F. van, Wolff, P., Martínez, G., Köhler, C., and Martienssen, R.A. (2018). Transposon-derived small RNAs triggered by miR845 mediate genome dosage response in Arabidopsis. Nat. Genet. 50, 186.

Burkart-Waco, D., Ngo, K., Dilkes, B., Josefsson, C., and Comai, L. (2013). Early Disruption of Maternal–Zygotic Interaction and Activation of Defense-Like Responses in Arabidopsis Interspecific Crosses. Plant Cell 25, 2037–2055.

Castillo, D.M., and Moyle, L.C. (2012). Evolutionary Implications of Mechanistic Models of TE-Mediated Hybrid Incompatibility. Int J Evol Biol 2012, 698198

Cooper, D.C. (1951). Caryopsis Development Following Matings Between Diploid and Tetraploid Strains of Zea Mays. Am. J. Bot. 38, 702–708.

Creasey, K.M., Zhai, J., Borges, F., Van Ex, F., Regulski, M., Meyers, B.C., and Martienssen, R.A. (2014). miRNAs trigger widespread epigenetically activated siRNAs from transposons in *Arabidopsis*. Nature 508, 411–415.

Dilkes, B.P., Spielman, M., Weizbauer, R., Watson, B., Burkart-Waco, D., Scott, R.J., and Comai, L. (2008). The Maternally Expressed WRKY Transcription Factor TTG2 Controls Lethality in Interploidy Crosses of Arabidopsis. PLOS Biol. 6, e308.

Doughty, J., Aljabri, M., and Scott, R.J. (2014). Flavonoids and the regulation of seed size in Arabidopsis. Biochem. Soc. Trans. 42, 364–369.

Erdmann, R.M., Satyaki, P.R.V., Klosinska, M., and Gehring, M. (2017). A Small RNA Pathway Mediates Allelic Dosage in Endosperm. Cell Rep. 21, 3364–3372.

d’Erfurth, I., Jolivet, S., Froger, N., Catrice, O., Novatchkova, M., and Mercier, R. (2009). Turning Meiosis into Mitosis. PLOS Biol. 7, e1000124.

Esen, A., and Soost, R.K. (1973). Seed Development in Citrus with Special Reference to 2x X 4x Crosses. Am. J. Bot. 60, 448–462.

Figueiredo, D.D., Batista, R.A., Roszak, P.J., Hennig, L., and Köhler, C. (2016). Auxin production in the endosperm drives seed coat development in Arabidopsis. ELife 5, e20542.

Fiume, E., and Fletcher, J.C. (2012). Regulation of Arabidopsis Embryo and Endosperm Development by the Polypeptide Signaling Molecule CLE8. Plant Cell 24, 1000–1012.

Gasciolli, V., Mallory, A.C., Bartel, D.P., and Vaucheret, H. (2005). Partially redundant functions of Arabidopsis DICER-like enzymes and a role for DCL4 in producing trans-acting siRNAs. Curr. Biol. 15, 1494–1500.

Gehring, M., Bubb, K.L., and Henikoff, S. (2009). Extensive Demethylation of Repetitive Elements During Seed Development Underlies Gene Imprinting. Science 324, 1447–1451.

Gehring, M., Missirian, V., and Henikoff, S. (2011). Genomic analysis of parent-of-origin allelic expression in Arabidopsis thaliana seeds. PloS One 6, e23687.

Gutierrez-Marcos, J.F., Pennington, P.D., Costa, L.M., and Dickinson, H.G. (2003). Imprinting in the endosperm: a possible role in preventing wide hybridization. Philos. Trans. R. Soc. Lond. B Biol. Sci. 358, 1105–1111.

Haig, and Westboy (1991). Genomic imprinting in endosperm: its effect on seed development in crosses between species, and its implications for the evolution of apomixis. Philos. Trans. R. Soc. Lond. B. Biol. Sci. 333, 1–13.

Håkansson, A., and Ellerström, S. (1950). SEED DEVELOPMENT AFTER RECIPROCAL CROSSES BETWEEN DIPLOID AND TETRAPLOID RYE. Hereditas 36, 256–296.

Hehenberger, E., Kradolfer, D., and Köhler, C. (2012). Endosperm cellularization defines an important developmental transition for embryo development. Development 139, 2031–2039.

Hsieh, T.-F., Ibarra, C.A., Silva, P., Zemach, A., Eshed-Williams, L., Fischer, R.L., and Zilberman, D. (2009). Genome-Wide Demethylation of Arabidopsis Endosperm. Science 324, 1451–1454.

Huang, F., Zhu, Q.-H., Zhu, A., Wu, X., Xie, L., Wu, X., Helliwell, C., Chaudhury, A., Finnegan, E.J., and Luo, M. (2017). Mutants in the imprinted PICKLE RELATED 2 gene suppress seed abortion of fertilization independent seed class mutants and paternal excess interploidy crosses in Arabidopsis. Plant J. 90, 383–395.

Ibarra, C.A., Feng, X., Schoft, V.K., Hsieh, T.-F., Uzawa, R., Rodrigues, J.A., Zemach, A., Chumak, N., Machlicova, A., Nishimura, T., et al. (2012). Active DNA Demethylation in Plant Companion Cells Reinforces Transposon Methylation in Gametes. Science 337, 1360–1364.

Jiang, H., Moreno-Romero, J., Santos-González, J., Jaeger, G.D., Gevaert, K., Slijke, E.V.D., and Köhler, C. (2017). Ectopic application of the repressive histone modification H3K9me2 establishes post-zygotic reproductive isolation in Arabidopsis thaliana. Genes Dev. 31: 1272–1287

Jurka, J., Kapitonov, V.V., Pavlicek, A., Klonowski, P., Kohany, O., and Walichiewicz, J. (2005). Repbase Update, a database of eukaryotic repetitive elements. Cytogenet. Genome Res. 110, 462–467.

Kelleher, E.S. (2016). Reexamining the P-Element Invasion of Drosophila melanogaster Through the Lens of piRNA Silencing. Genetics 203, 1513–1531.

Kim, D., Pertea, G., Trapnell, C., Pimentel, H., Kelley, R., and Salzberg, S.L. (2013). TopHat2: accurate alignment of transcriptomes in the presence of insertions, deletions and gene fusions. Genome Biol. 14, R36.

Kradolfer, D., Wolff, P., Jiang, H., Siretskiy, A., and Köhler, C. (2013). An imprinted gene underlies postzygotic reproductive isolation in Arabidopsis thaliana. Dev. Cell 26, 525–535.

Lafon-Placette, C., and Köhler, C. (2015). Epigenetic mechanisms of postzygotic reproductive isolation in plants. Curr. Opin. Plant Biol. 23, 39–44.

Law, J.A., Du, J., Hale, C.J., Feng, S., Krajewski, K., Palanca, A.M.S., Strahl, B.D., Patel, D.J., and Jacobsen, S.E. (2013). Polymerase IV occupancy at RNA-directed DNA methylation sites requires SHH1. Nature 498, 385–389.

Li, J., and Berger, F. (2012). Endosperm: food for humankind and fodder for scientific discoveries. New Phytol. 195, 290–305.

Love, M.I., Huber, W., and Anders, S. (2014). Moderated estimation of fold change and dispersion for RNA-seq data with DESeq2. Genome Biol. 15, 550.

Lu, J., Zhang, C., Baulcombe, D.C., and Chen, Z.J. (2012). Maternal siRNAs as regulators of parental genome imbalance and gene expression in endosperm of Arabidopsis seeds. Proc. Natl. Acad. Sci. U. S. A. 109, 5529–5534.

Marí-Ordóñez, A., Marchais, A., Etcheverry, M., Martin, A., Colot, V., and Voinnet, O. (2013). Reconstructing *de novo* silencing of an active plant retrotransposon. Nat. Genet. 45, 1029–1039.

Martienssen, R.A. (2010). Heterochromatin, small RNA and post-fertilization dysgenesis in allopolyploid and interploid hybrids of Arabidopsis. New Phytol. 186, 46–53.

Martinez, G., Wolff, P., Wang, Z., Moreno-Romero, J., Santos-González, J., Conze, L.L., DeFraia, C., Slotkin, R.K., and Köhler, C. (2018). Paternal easiRNAs regulate parental genome dosage in Arabidopsis. Nat. Genet. 50, 193.

McCue, A.D., Nuthikattu, S., Reeder, S.H., and Slotkin, R.K. (2012). Gene Expression and Stress Response Mediated by the Epigenetic Regulation of a Transposable Element Small RNA. PLOS Genet. 8, e1002474.

Milbocker, D., and Sink, K. (1969). Embryology of diploid × diploid and diploid × tetraploid crosses in poinsettia. Can. J. Genet. Cytol. 11, 598–601.

Mittelsten Scheid, O., Afsar, K., and Paszkowski, J. (2003). Formation of stable epialleles and their paramutation-like interaction in tetraploid *Arabidopsis thaliana*. Nat. Genet. 34, 450–454.

Muntzing, A. (1936). The Evolutionary Significance of Autopolyploidy. Hereditas 21: 363–378

Oberlin, S., Sarazin, A., Chevalier, C., Voinnet, O., and Marí-Ordóñez, A. (2017). A genome-wide transcriptome and translatome analysis of Arabidopsis transposons identifies a unique and conserved genome expression strategy for Ty1/Copia retroelements. Genome Res. 27, 1549–1562.

Pignatta, D., Erdmann, R.M., Scheer, E., Picard, C.L., Bell, G.W., and Gehring, M. (2014). Natural epigenetic polymorphisms lead to intraspecific variation in Arabidopsis gene imprinting. ELife 3, e03198.

Pignatta, D., Bell, G., and Gehring, M. (2015). Whole Genome Bisulfite Sequencing and DNA Methylation Analysis from Plant Tissue. BIO-Protoc. 5.

Piskurewicz, U., Iwasaki, M., Susaki, D., Megies, C., Kinoshita, T., and Lopez-Molina, L. (2016). Dormancy-specific imprinting underlies maternal inheritance of seed dormancy in Arabidopsis thaliana. ELife 5, e19573.

Povilus, R.A., Diggle, P.K., and Friedman, W.E. (2018). Evidence for parent-of-origin effects and interparental conflict in seeds of an ancient flowering plant lineage. Proc. Biol. Sci. 285.

Quinlan, A.R., and Hall, I.M. (2010). BEDTools: a flexible suite of utilities for comparing genomic features. Bioinformatics 26, 841–842.

Satyaki, P.R.V., and Gehring, M. (2017). DNA methylation and imprinting in plants: machinery and mechanisms. Crit. Rev. Biochem. Mol. Biol. 52, 163–175.

Schatlowski, N., Wolff, P., Santos-González, J., Schoft, V., Siretskiy, A., Scott, R., Tamaru, H., and Köhler, C. (2014). Hypomethylated Pollen Bypasses the Interploidy Hybridization Barrier in Arabidopsis. Plant Cell 26, 3556–3568.

Schon, M.A., and Nodine, M.D. (2017). Widespread Contamination of Arabidopsis Embryo and Endosperm Transcriptome Data Sets. Plant Cell 29, 608–617.

Scott, R.J., Spielman, M., Bailey, J., and Dickinson, H.G. (1998). Parent-of-origin effects on seed development in Arabidopsis thaliana. Development 125, 3329–3341.

Stoute, A.I., Varenko, V., King, G.J., Scott, R.J., and Kurup, S. (2012). Parental genome imbalance in Brassica oleracea causes asymmetric triploid block: Parent-of-origin effects in Brassica oleracea. Plant J. 71: 503–516.

Stroud, H., Greenberg, M.V.C., Feng, S., Bernatavichute, Y.V., and Jacobsen, S.E. (2013). Comprehensive Analysis of Silencing Mutants Reveals Complex Regulation of the Arabidopsis Methylome. Cell 152, 352–364.

Tiwari, S., Spielman, M., Schulz, R., Oakey, R.J., Kelsey, G., Salazar, A., Zhang, K., Pennell, R., and Scott, R.J. (2010). Transcriptional profiles underlying parent-of-origin effects in seeds of Arabidopsis thaliana. BMC Plant Biol. 10, 72.

Tsay, Y.F., Frank, M.J., Page, T., Dean, C., and Crawford, N.M. (1993). Identification of a mobile endogenous transposon in Arabidopsis thaliana. Science 260, 342–344.

Vu, T.M., Nakamura, M., Calarco, J.P., Susaki, D., Lim, P.Q., Kinoshita, T., Higashiyama, T., Martienssen, R.A., and Berger, F. (2013). RNA-directed DNA methylation regulates parental genomic imprinting at several loci in Arabidopsis. Development 140, 2953–2960.

Walia, H., Josefsson, C., Dilkes, B., Kirkbride, R., Harada, J., and Comai, L. (2009). Dosage-Dependent Deregulation of an AGAMOUS-LIKE Gene Cluster Contributes to Interspecific Incompatibility. Curr. Biol. 19, 1128–1132.

Walker, J., Gao, H., Zhang, J., Aldridge, B., Vickers, M., Higgins, J.D., and Feng, X. (2018). Sexual-lineage-specific DNA methylation regulates meiosis in Arabidopsis. Nat. Genet. 50, 130.

Wang, G., Jiang, H., Del Toro de León, G., Martinez, G., and Köhler, C. (2018). Sequestration of a Transposon-Derived siRNA by a Target Mimic Imprinted Gene Induces Postzygotic Reproductive Isolation in Arabidopsis. Dev. Cell 46, 696–705.

Wang, L., Feng, Z., Wang, X., Wang, X., and Zhang, X. (2010). DEGseq: an R package for identifying differentially expressed genes from RNA-seq data. Bioinforma. Oxf. Engl. 26, 136–138.

von Wangenheim, K.-H., and Peterson, H.-P. (2004). Aberrant endosperm development in interploidy crosses reveals a timer of differentiation. Dev. Biol. 270, 277–289.

Williams, B.P., Pignatta, D., Henikoff, S., and Gehring, M. (2015). Methylation-Sensitive Expression of a DNA Demethylase Gene Serves As an Epigenetic Rheostat. PLOS Genet. 11, e1005142.

Wolff, P., Jiang, H., Wang, G., Santos-González, J., and Köhler, C. (2015). Paternally expressed imprinted genes establish postzygotic hybridization barriers in Arabidopsis thaliana. ELife 4:e10074.

Xiao, W., Brown, R.C., Lemmon, B.E., Harada, J.J., Goldberg, R.B., and Fischer, R.L. (2006). Regulation of Seed Size by Hypomethylation of Maternal and Paternal Genomes. Plant Physiol. 142, 1160–1168.

Yan, D., Duermeyer, L., Leoveanu, C., and Nambara, E. (2014). The Functions of the Endosperm During Seed Germination. Plant Cell Physiol. 55, 1521–1533.

Yu, Z., Haberer, G., Matthes, M., Rattei, T., Mayer, K.F.X., Gierl, A., and Torres-Ruiz, R.A. (2010). Impact of natural genetic variation on the transcriptome of autotetraploid Arabidopsis thaliana. Proc. Natl. Acad. Sci. 107, 17809–17814.

Zhang, J., Liu, Y., Xia, E.-H., Yao, Q.-Y., Liu, X.-D., and Gao, L.-Z. (2015). Autotetraploid rice methylome analysis reveals methylation variation of transposable elements and their effects on gene expression. Proc. Natl. Acad. Sci. U. S. A. 112, E7022–7029.

Zhang, Y., Malone, J.H., Powell, S.K., Periwal, V., Spana, E., MacAlpine, D.M., and Oliver, B. (2010). Expression in Aneuploid Drosophila S2 Cells. PLOS Biol. 8, e1000320.

Zilberman, D., Gehring, M., Tran, R.K., Ballinger, T., and Henikoff, S. (2007). Genome-wide analysis of *Arabidopsis thaliana* DNA methylation uncovers an interdependence between methylation and transcription. Nat. Genet. 39, 61–69.

